# Tracking cellular biomolecular condensate dynamics under proteostatic stress with middle-down phosphoproteomics

**DOI:** 10.64898/2026.09.22.753649

**Authors:** Boomathi Pandi, Dominic C.M. Ng, Peyton Schaal, Lorena Alamillo, Shaonil Binti, Edward Lau, Maggie P.Y. Lam

**Affiliations:** Department of Medicine; Department of Biochemistry & Molecular Genetics University of Colorado School of Medicine Aurora, CO 80045, USA

**Keywords:** Biomolecular condensates, splice factors, partition, solubility, middle-down proteomics

## Abstract

Coordination of biological function requires the partition of cellular components including into biomolecular condensates, but an overall landscape of how protein compartmentalize into higher-order assemblies under stress is still emerging. We apply proteome-wide solubility profiling to compare the compositions of NP-40-insoluble proteins and their phosphorylation status, using a new mass spectrometry-based hybrid bottom-up and chemical middle-down proteomics approach to analyze the solubility behavior of 8,740 proteins and 31,647 phosphopeptides under normal and ER stress conditions. Cell stress induces a pervasive differential partition of proteins in and out of detergent-insoluble cellular compartments. This differential partition is partially orthogonal to stress-induced abundance changes and comprises both phosphorylation-dependent and phosphorylation-independent mechanisms. Whereas phosphorylation-independent partition changes involve largely secretory pathway proteins and implicate higher-order assemblies of chaperones and clients, phosphorylation-based partitions suggest a dynamic rearrangement of biomolecular condensate compositions across cytoplasmic and nuclear ribonucleoprotein assemblies. The accumulation of serine/arginine rich (SR) proteins and other annotated nuclear speckle members in the condensate-rich proteome fractions emerges as a central feature of stress-induced remodeling. Our results establish global solubility dynamics as an integral component of proteome stress response and implicates broad involvements of splice factor spatial reorganization as a prominent facet of ER stress response.

## Introduction

ER stress is commonly associated with multiple human diseases. During unfolded protein response (UPR) following proteostatic stress, cytoplasmic ribonucleoprotein granules form to sequester ribosome and ribonucleoprotein complexes, which are thought to protect transcripts from degradation under global translation arrest. Recent work has uncovered multiple aspects of UPR signaling than previously appreciated, including the synthesis of splice factors leading to concerted changes in splicing programs (1) and suggesting a complex network of multifaceted cell wide response to proteostatic stress. Biomolecular condensates are a fundamental mode of spatial organization in the cell. The selective partition of molecules into condensates underpins the formation and dynamics of membraneless organelles, such as the nucleolus, nuclear speckle, and stress granules, and is known to play critical roles in transcriptional and translational regulations (2,3). A key distinction of membraneless assemblies over membraned organelles is that they can dynamically form and dissolve in response to cellular and environmental signals (4,5). This places membraneless organelles including the nucleolus and the nuclear speckles as dynamic centers that coordinate cellular stress response (2,6,7). However, although much progress has been made on the organization principles of individual condensates such as stress granules and nucleolus, a proteome-wide view is missing on the dynamic partition of proteins into or out of biomolecular condensates and their functional significance during stress response.

Proteins that participate in biomolecular condensation of ribonucleoprotein granules, including the serine/arginine-rich splice factor (SRSF) and heterogeneous nuclear ribonucleoprotein (hnRNP) families of RNA-binding proteins (RBPs), are enriched in intrinsically disordered, low-complexity sequences (8–10). Protein intrinsically disordered domains (IDR) play pivotal roles in multivalent molecular interactions and phase separation behaviors, and are marked by unique preferences in sequence compositions, including low complexity domains and charge blocks (8–12). These sequences frequently contain high lysine and arginine density, making them unamenable to producing suitable tryptic peptides, and moreover are enriched in accessible serine and threonine that make them protein phosphorylation hotspots.

By altering the charge patterns of IDR, phosphorylation is well placed as a mechanism to tune the strength of charge-based multivalent interactions, such as between proteins or between protein and RNA, dynamically upon cellular and environmental cues. For instance, the phosphorylation status of RNA polymerase II drives its distribution across biomolecular condensates, whereas the phosphorylation of SRSF proteins and other RBPs are known to control their nucleocytoplasmic shuttling and biomolecular condensate formation (13–17). While it is known that certain stress conditions including heat shock and hypoxia can alter the phosphorylation of SRSF proteins (17–20), a systematic picture is still emerging on the phosphorylation profiles of condensate-forming proteins upon cell stress, and their effects on proteome organization.

Here we characterized the dynamic partition of proteins and phosphoproteins into detergent soluble and insoluble fractions under prolonged ER stress, with the goal of mapping the global remodeling of membraneless organelles and other higher-order assemblies via the formation of biomolecular condensates. Our strategy employs innovative protein and phosphoprotein solubility profiling coupled to liquid chromatography and tandem mass spectrometry (LC-MS/MS) to identify the global solubility behaviors of proteins in normal and stressed cells. The findings reveal broad remodeling of protein solubility behaviors in stressed cells, involving thousands of proteoforms associated with both cytoplasmic and nuclear bodies including stress granules, nucleoli, nuclear speckles, and transcriptional condensates. Protein partition behaviors can be categorized as phosphorylation-dependent and phosphorylation-independent regulations. The partition of RBP involved in nuclear speckles and splicing regulations emerges as a prominent feature of prolonged ER stress. Together, our results broaden our current understanding of the ER stress and UPR regulon, revealing a proteome-wide organization principle of RBPs relevant to stress response splicing program. The breadth of protein re-distribution behaviors, often in the absence of detectable abundance changes, may have implications for the discovery of intervention targets in ER stress found in a variety of diseases.

## Results

### Proteome-wide profiling of protein solubility with hybrid bottom-up middle-down proteomics

We employ an optimized mass spectrometry-based solubility proteomics method to find the differential partition of proteins into soluble and insoluble fractions under stress. Mild detergents such as NP-40 can be used to biochemically enrich for higher-order protein and protein-RNA assemblies (21–23). Upon NP- 40 solubilization and centrifugation, membranes are disrupted and soluble proteins are released to the supernatant, leaving insoluble proteins from biomolecular condensates and other higher-order assemblies within the pellet fraction. Here, by focusing on deep characterization of the NP-40-insoluble pellets using a combination of bottom-up and middle-down proteomics, we directly analyze the proteins that are selectively concentrated within biomolecular condensates and their quantitative changes in stress, which may escape detection in the soluble fraction (**Figure 1A**). By incorporating a TiO_2_/Fe-NTA double phosphopeptide enrichment, we further enable deep profiling of phosphorylation sites on both acidic and basic phosphopeptides.

**Figure 1:**
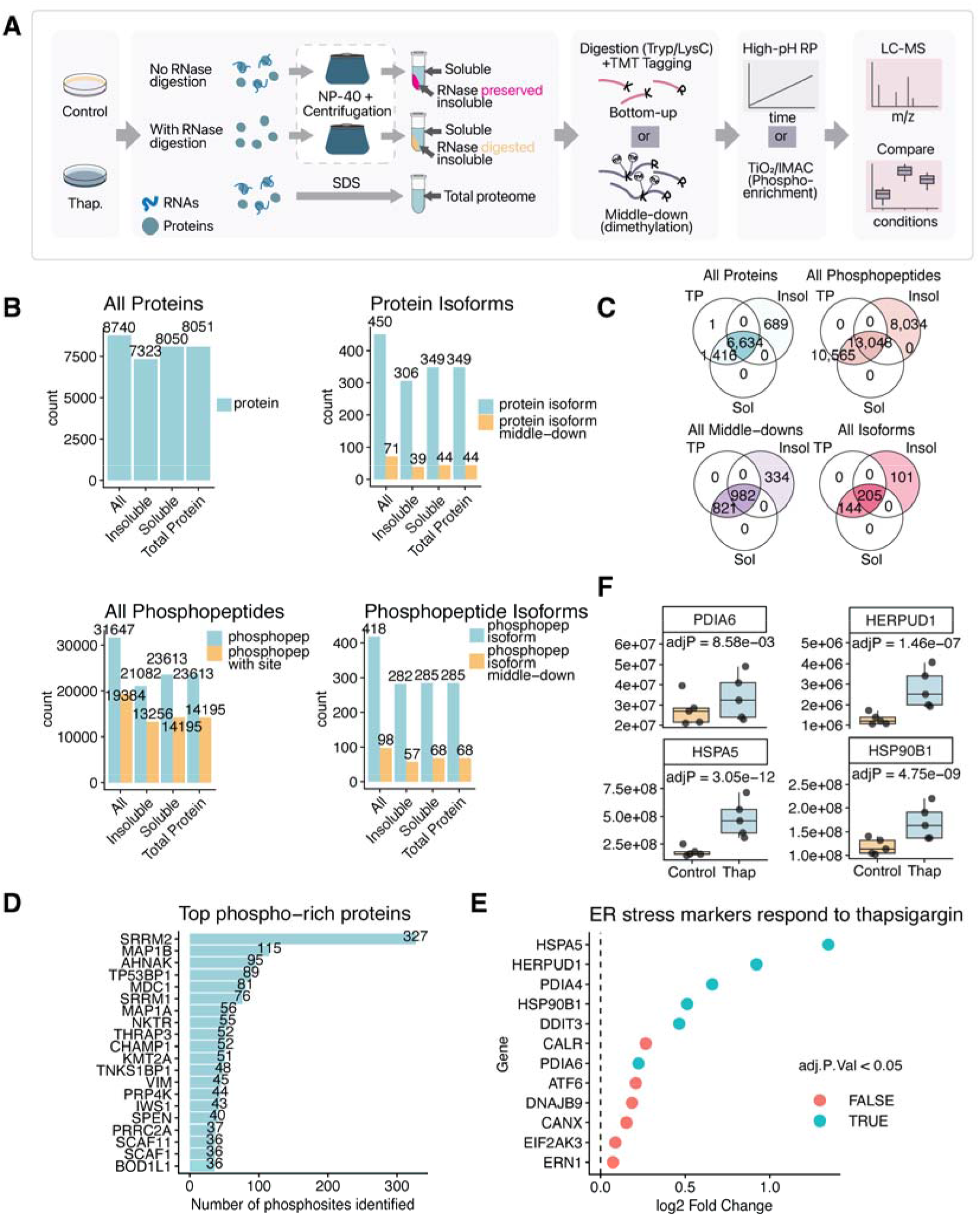
Experimental design of solubility proteomics and phosphoproteomics experiments. A. Workflow of hybrid middle-down/bottom-up proteomics approach to measure stress-induced protein differential solubility. AC16 cells are treated with or without thapsigargin. Extracted cells are either solubilized with NP-40 (with or without RNase digestion) or SDS. NP-40 treated samples undergoes ultracentrifugation to separate proteins into soluble and insoluble fractions prior to chemical dimethylation for middle-down TMT proteomics or bottom-up proteomics without dimethylation. Phosphopeptide enrichments are performed in parallel with TiO2/IMAC columns prior to TMT-MS. B. Number of proteins identified in each condition. From top left clockwise: All proteins identified in soluble and insoluble fractions; number of non-canonical (alternative) protein isoforms identified in total, and those identified only via middle-down proteomics; all phosphopeptides identified and phosphopeptides identified with unambiguous site assignment (PTM-Prophet); all phosphopetpides belonging to alternative protein isoforms and those identified only via middle-down proteomics. C. Top: Venn diagrams showing the overlap of proteins (left) and phosphopeptides (right) identified in each of the total protein (TP), insoluble (insol), and soluble (sol) fractions and their overlaps. Bottom: Venn diagrams showing the overlaps in proteins identified only via middle-down (left) and all identified alternative protein isoforms (right) in each fraction as above. D. Top 20 proteins with most number of phosphorylation site coverage in the experiment. E. Induction of known ER stress markers upon thapsigargin treatment. Color: limma adjusted P values < 0.05. F. Individual boxplots of differentially expressed ER stress marker proteins PDIA6, HERPUD1, HSPA5, and HSP90B1.

We applied this method to normal and ER stressed cells over 5 biological replicates, followed by differential solubility and phosphorylation enrichments. The resulting insoluble and soluble fractions were separately labeled tandem mass tags (TMT) quantitative proteomics experiments, analyzing a total of 8740 distinct proteins, including 450 non-canonical protein isoforms and 31,647 distinct phosphopeptides that can be confidently assigned to 19,384 phosphorylation sites (**Figure 1B**). The TiO2/Fe-NTA enrichment identified multiple phosphoproteins with a large number of phosphorylation sites, with examples including SRRM2 with 327 phosphorylation sites and MAP1B with 115 phosphorylation sites (**Figure 1D**).

As expected, prolonged ER stress induced by 16 hours of thapsigargin treatment led to significant up- regulations of protein-level ER stress markers (**Figure 1E**), including significant upregulation of HSPA5/BiP/Grp78, HERPUD1, HSP90B1/Grp94, and PDIA6 (**Figure 1F**), reflecting a robust stress induction upon thapsigargin treatment. To further confirm the tested dosage and time regime is relevant to stress-induced condensates, we expressed a G3BP1-SNAP construct and confirmed puncta formation under thapsigargin-treated but not untreated cells; moreover, the degree of puncta formation under stress varies in G3BP1-S149A and S149E constructs suggesting it is in part regulated by the known G3BP1 RNA-binding phosphoswitch at the S149 site (**Supplementary Figure S1**). Therefore, we achieved deep proteome coverage to allow comparison of their solubility behaviors between normal and stressed cells in a manner relevant to prolonged UPR and ER stress response.

The chemical middle-down approach contributed additional peptides, proteins, and phosphosites not seen in the bottom-up experiments. To enable middle-down coverage, we chemically dimethylated intact proteins to block free internal lysines, followed by trypsin cleavage of peptide bonds only following arginines (**Figure 2A**). This allows the potential to improve protein coverage, similar to the commonly-used middle-down protease LysC, while retaining the robustness, site-specificity, and low-cost of trypsin digestion. To determine whether the increased coverage was not simply due to stochastic non-overlaps of repeat shotgun proteomics profiling experiments, we examined the physiochemical properties of identified peptide in each digestion method. Dimethylation-based middle-down identified peptides with significantly greater average lengths, and significantly greater proportion of peptides with at least 20 (34.8% vs. 15.3%) or 30 (10.2% vs. 3.3%) residues over bottom-up coverage (**Figure 2B**). The dimethylation protection is near-complete as virtually all identified peptides have internal miscleaved lysines (**Figure 2C**). The identified middle-down peptides overlap minimally with the bottom-up experiments in sequence (**Figure 2D**, bars 1–3 vs. bars 4–6). Moreover, when we consider the fully-tryptic fragments of the middle-down peptides (i.e., cleaved at the dimethylated lysines), fewer than half of the middle-down detected peptides contained internal fragments identified in the bottom-up experiments (**Figure 2D**, bars 1–2 vs. bar 3). Importantly, a considerable number of middle-down peptides are composed only of full-tryptic fragments that are less than 6 residues long each (**Figure 2D**, bar 2) which would render them operationally undetectable in typical proteomics experiment and indicate a mechanism for middle-down proteomics to increase coverage even from peptides that are otherwise within the normal length distributions of a bottom-up experiment.

**Figure 2:**
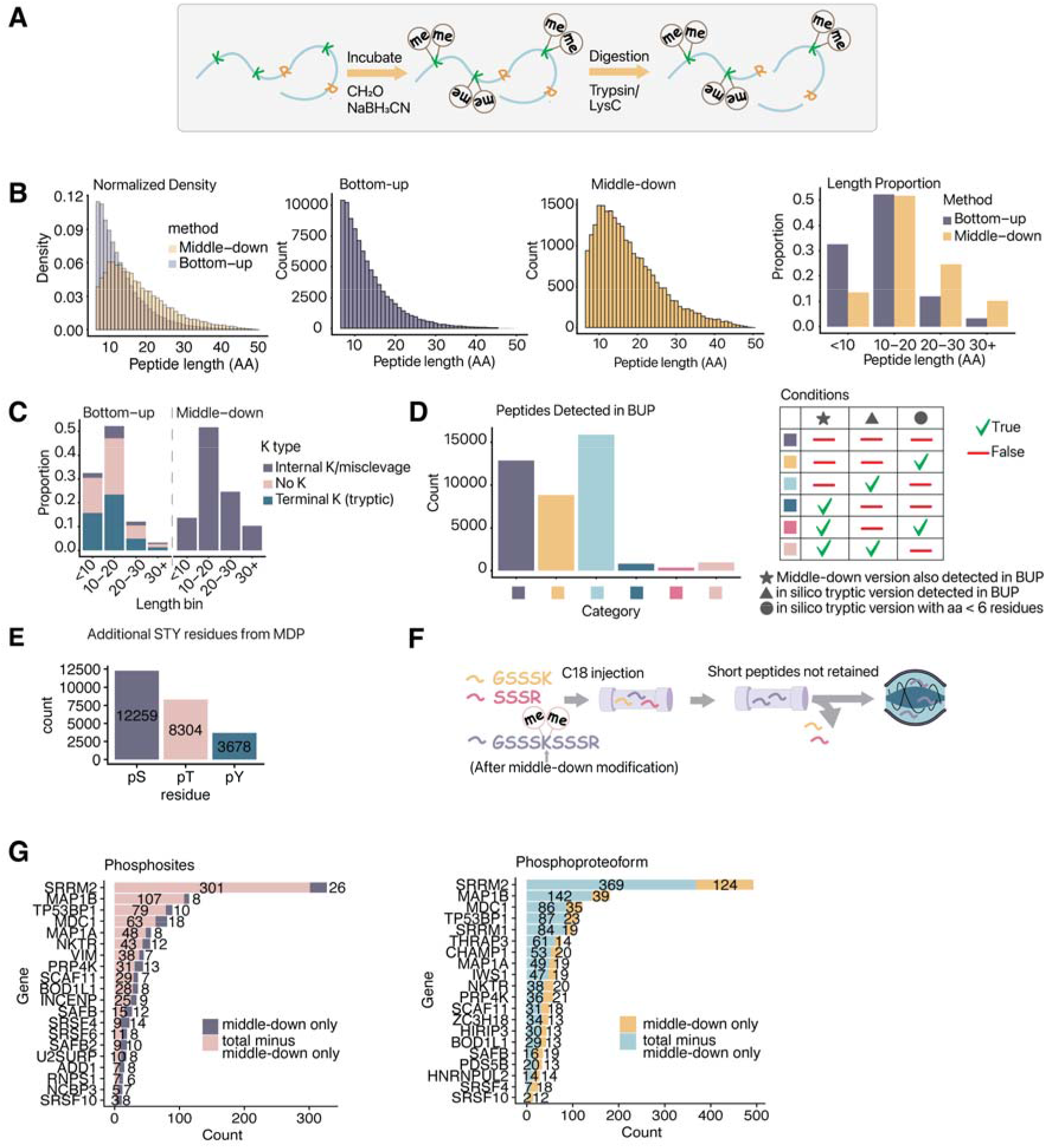
Middle-down proteomics via chemical demethylation increases IDR and proteoform coverage. A. Schematics of dimethylation of intact proteins prior to digestion for middle-down peptide generation. B. From left to right: count-normalized density of peptide length distributions for peptides identified from middle-down (orange), and bottom-up (grey) experiments; peptide length distributions of bottom-up experiments; peptide length distribution of middle-down experiments; proportion of peptides with length across bins. C. Proportional bar charts comparing the number of peptides with internal and terminal lysine (K) residues in bottom-up and middle-down proteomics experiments. D. Among peptides detected in middle-down experiments, we consider the number of these peptides for which an identical sequence is identified in bottom-up experiments. Second, we consider if we digest the middle-down peptides in silico by trypsin rule, i.e., cleaving the internal K, whether these peptides are detected in bottom-up proteomics. Finally, we consider whether the in-silico digested internal tryptic peptides are less than 6 amino acids long, which would render them unlikely to be detectable in LC-MS/MS experiments. The number of cases from the combinations of these scenarios are shown. E. Bar charts showing the number of additional phosphorylatable (serine/S; threonine/T; and tyrosine/Y) residues detected only via the middle-down approach in the data. F. Conceptual figure depicting the concept that middle-down proteomics retain peptides that are otherwise too short to be detectable in bottom-up proteomics experiment. G. Middle-down proteomics enables greater numbers of phosphorylation sites (left) and phosphoproteoforms among splicing-related RNA-binding proteins.

In parallel, the middle-down approach added significantly to the number of serine (12259 extra residues covered), threonine (8304 residues), and tyrosine (3678 residues), indicating its potential to add additional phosphorylation sites to the study coverage (**Figure 2E**). This has an outsized effects on short, intrinsically disordered proteins including SR and hnRNP proteins. For instance, although middle-down proteomics adds only a minor proportion of phosphorylation sites to all the sites identified by bottom-up proteomics, in certain proteins including SRSF4 (14/23 sites), SRSF6 (8/19 sites), and SRSF10 (8/11 sites), a substantial portion of identified sites were contributed only by middle-down proteomics (**Figure 2G**); whereas for HNRNPA1, HNRNPH1, and HNRNPF, all identified sites were from middle-down proteomics (graph not showing this currently because of how we selected/ordered sites). Middle-down peptides also contributed substantially to distinct phosphopeptides, including those harboring unique combinations of one or more phosphorylation sites, i.e., phosphoproteoforms; for instance, in SRRM2, whereas only 26 out of 327 phosphorylation sites (8%) are identified through middle-down proteomics only, 124 out of 493 phosphoproteoforms (25%) are contributed by middle-down peptides (**Figure 2G**; right). Taken together, this analysis show that our chemical middle-down approach adds significantly to protein quantification coverage and may take on further importance in lysine-rich, phosphorylation-rich IDRs such as those enriched in biomolecular condensates, where a regular trypsin digestion may result in peptides that are too short to support confident identification (**Figure 2F**). These results support our rationale of using a hybrid bottom-up middle-down approach to characterize protein solubility partition in stressed cells.

### ER stress induces phosphorylation dependent and independent protein differential partition

To distinguish various modalities of proteomic responses during stress, we consider several scenarios of proteome reconfigurations that may be detected in the experiment (**Figure 3A**). Upon thapsigargin-induced ER stress, a protein may alter in abundance in the total proteome, i.e., differential expression (**Figure 3A**, Box 1). In parallel, a protein pool may selectively partition into insoluble fractions, in the absence of total abundance change, suggesting a redistribution of proteins (Box 2). In a more general scenario, a protein may also change in total abundance, but regardless, the selective partition behavior will manifest in a different stressed-to-normal abundance ratio between the insoluble and the total protein pool, indicating a change in partition coefficients (Box 3). Moreover, a protein may be detectable only within the insoluble fraction due to the enrichment effect, i.e., only a subset of the proteome is found insoluble, rendering them more detectable (Box 4). These scenarios also intersect with the phosphorylation status of the protein, such that the phosphoproteoform may exhibit a different partition behavior than the native (unmodified) counterpart (Box 5).

**Figure 3:**
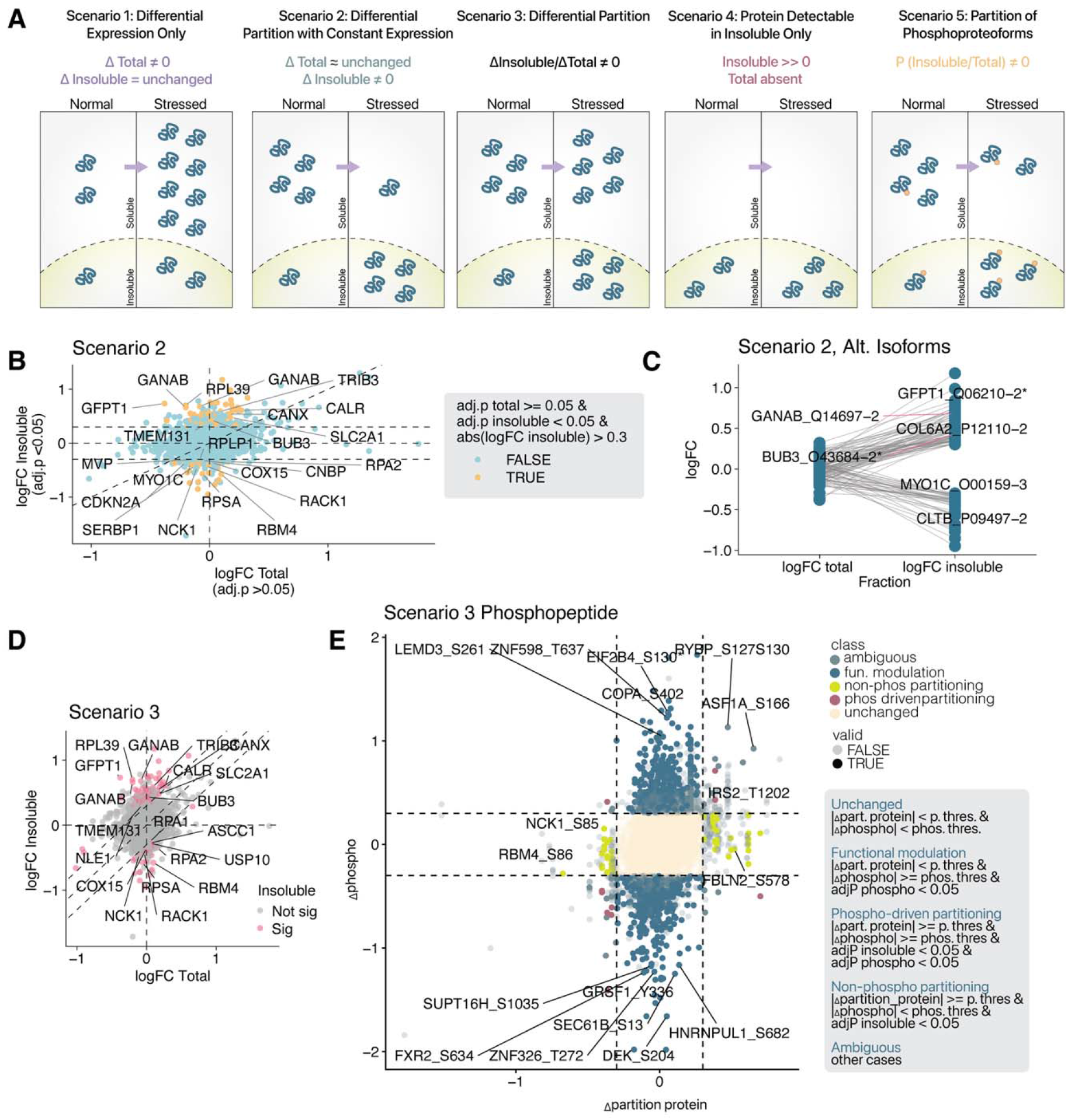
ER stress induces proteome-wide alterations in protein solubility partition. A. Several scenarios outlining differential partition between normal and stressed (thapsigargin-treated) cells are illustrated. Scenario 1 (**S1**): A protein exhibits differential expression only (i.e., Δ total protein ≠ 0) but minimal differences in the insoluble fraction or the insoluble to total ratio. **S2:** A protein exhibits differential partition in the absence of overt different expression, i.e., limma adj. P (thapsigargin/control) in total protein fraction ≥ 0.05 while limma adj. P (thapsigargin/control) < 0.05 and |logFC| > 0.3 in the insoluble fraction. **S3.** A more general case is a partition regardless of whether the protein expression has changed, where Δ partition, defined as logFC (thapsigargin/control) in the insoluble fraction – logFC (thapsigargin/control) in the total proteome. **S4.** Because the NP-40 insoluble pellets are highly enriched in a subset of the proteome (i.e., aggregate and condensate-forming proteins), a protein may be detectable only through direct profiling of the insoluble pellets. **S5.** As an extended case of differential partitioning, a phosphoproteoform may become differentially distributed into the insoluble fraction in stressed cells independent of other phosphoproteoforms of the same protein, or its unmodified form. B. Scatterplot highlighting proteins with differential partition in the absence of differential expression in stressed cells (i.e., **S2**) (orange). The redistributed/partitioned proteins have limma adj. P (thapsigargin/control) ≥ 0.05 in the total proteins but limma adj. P < 0.05 and |logFC| > 0.3 in the insoluble fraction C. Per-protein differences in logFC (thapsigargin/control) between the total fraction and insoluble fraction of select proteins. The behavior of several non-canonical protein isoforms with differential partition (limma adj. P (thapsigargin/control) ≥ 0.05 in total proteins limma adj. P (thapsigargin/control) < 0.05 and |logFC| > 0.3 in the insoluble fraction) are shown. D. Scatterplot highlighting proteins with differential partition in stressed cells (i.e., **S3**). Proteins with limma adj. P (thapsigargin/control) < 0.05 in |Δ partition| >0.3 are highlighted in red. E. Scatterplots showing phosphopeptides with differential phosphorylation vs. the |Δ partition| of their unmodified protein counterpart, as a limited case of **S5**. Colors denote classes of interpreted behaviors based on a phosphopeptide’s differential phosphorylation and protein partition.

Focusing first on strict partition changes (Scenario 2), we discovered 197 proteins that show significant differences under stress only within NP-40-insoluble pellets (limma FDR adjusted P (adj. P) ≤ 0.05) but not in the total proteome (limma adj. P > 0.05), indicating broad redistribution of protein behavior is a veritable facet of prolonged ER stress orthogonal to differential protein expression. Whereas differential expression analysis turns up well established late-phase ER stress proteome markers including BiP/HSPA5, HYOU1, and HSP90B1, the differential partition comparison highlights GANAB (glucosidase II alpha), which is involved in protein synthesis and folding through removal of glucose from nascent glycoproteins in the ER; CANX/CALR, glycan-binding ER chaperones; TMEM131, a Golgi transport protein involved in collagen secretion; and BUB3, involved in mitotic checkpoints (**Figure 3B**, left). Moreover, these proteins encompass a number of poorly characterized alternative isoforms, including in particular some that are only specifically identifiable by middle-down peptides, such as BUB3-2, GFPT1- 2, and others; indicating solubility redistribution in the absence of differential expression impacts broad swathes of the proteome (**Figure 3C**). In some instances, isoforms of the same protein show differential regulations under ER stress, with the -2 isoform of EEF1D significantly decreased but not the -3 isoform.

We next considered a more general scenario where a protein shows differential partition coefficients into insoluble fractions under stress (Scenario 3), we discovered 241 proteins with differential levels in the NP-40 insoluble pellets (FDR adjusted P ≤ 0.05) regardless of whether their total protein level changes under ER stress (**Figure 3D**, left). Among these proteins are known ER-stress regulated proteins such as HSPA5 and HYOU1. As another example, XBP1s is an UPR protein known to oscillate under prolonged ER stress. We identify a peptide with pS288 corresponding to spliced XBP1s protein, which is up- regulated at the total protein level in 16 hours of thapsigargin over control cells. Interestingly, this behavior is more exclusive to the NP-40 insoluble pellet but not the soluble fraction, suggesting the elevated XBP1s is partitioned within insoluble cellular fractions (**Figure 4E**). Therefore, some stress response proteins may show altered biophysical behaviors (i.e., solubility partition) under stress in addition to their up-regulated expression. Other differentially partitioned proteins include BUB3 and GANAB and span diverse pathways, corroborating broad re-organization of the proteome.

**Figure 4:**
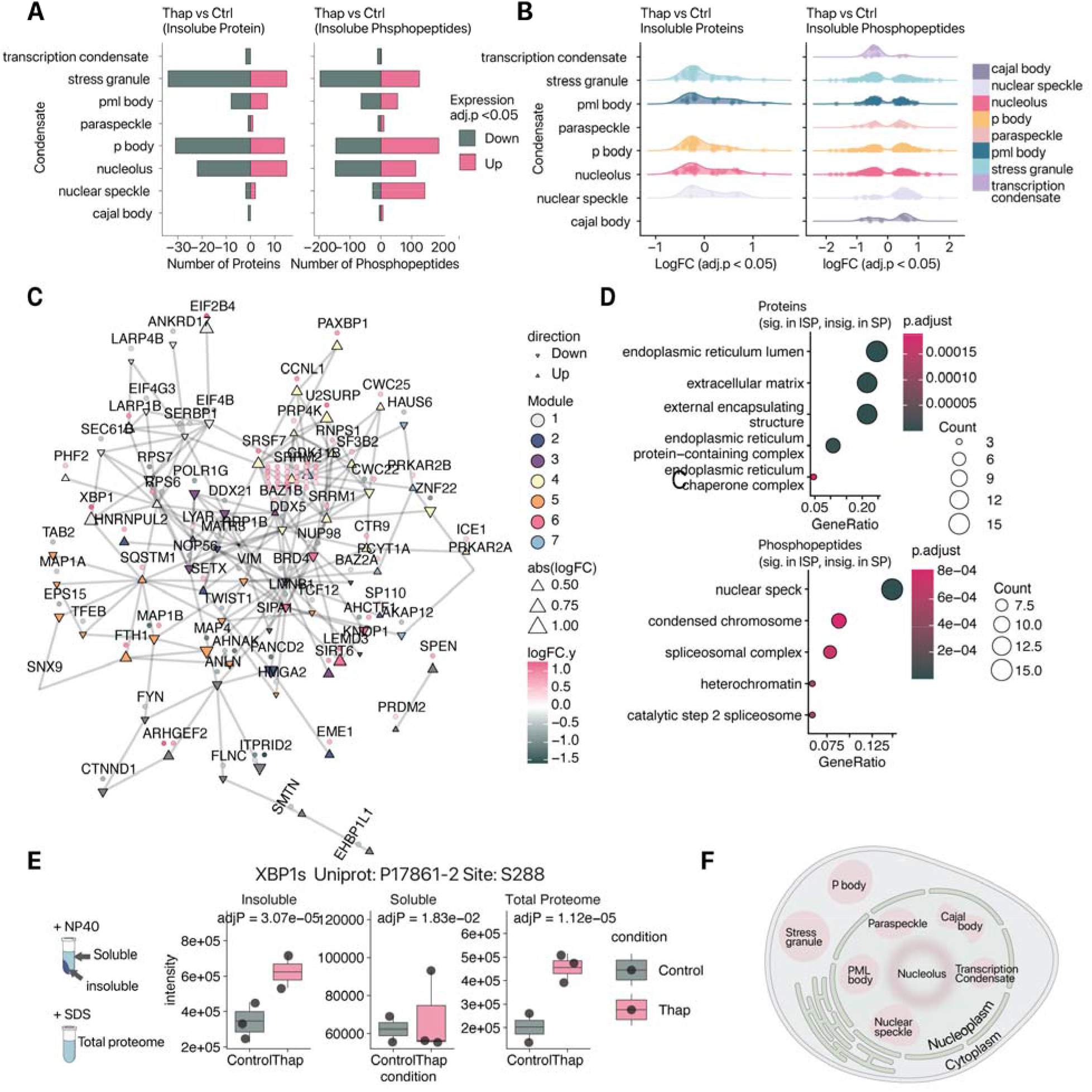
Insoluble proteome profiling reveals broad membraneless organelle remodeling. A. Stress-induced differential abundance (thapsigargin (Thap)/control (Ctrl) within the insoluble fraction) of proteins (left) and phosphopeptides of proteins (right) associated with various types of biomolecular condensate as annotated in CD-CODE. x-axis: number of proteins or phosphopeptides showing increased or decreased abundance; limma adj. P value < 0.05. B. As in A, but presented in ridge plots showing density of data points spread over range of logFC of each annotated condensate type in the insoluble fraction of thapsigargin (Thap) vs control (Ctrl) comparison; limma adj. P value < 0.05. C. Network graph showing the protein-protein interactions (edges) associated with quantified phosphopeptides, retrieved from StringDB. Proteins are included if they contain quantified phosphopeptides that are significantly altered within the insoluble fraction (thapsigargin/control; limma adj P < 0.01) but are not significantly altered in the soluble fraction (thapsigargin/control; limma adj P ≥ 0.01). Nodes shape represent direction of averaged logFC of all significantly changed phosphopeptide with site(s) identified (i.e., up or down), with the size of shape scaling with |logFC|. The individual logFC of each phosphopeptide associated with each protein is shown next to the nodes as filled circles. The Louvain algorithm is used to cluster and generate several modules (depicted with different node colors) based on network neighborhoods. Modules 2 contains proteins whose phosphopeptides are primarily repressed in the insoluble fraction, whereas Module 3, while module 3, containing SRRM2 and enriched in nuclear speckles proteins, are associated with up-regulated phosphopeptides. Thus, the solubility partition data reflects interconnected modules of proteins with coordinated behaviors. D. Top: GO overrepresentation analysis of unmodified proteins with significant differential abundance in the insoluble fraction (thapsigargin/control; limma adj. P < 0.01) but are not significantly changed in the soluble fraction (limma adj. P ≥ 0.01). Bottom: as in Top, but for phosphopeptides. E. The expression of the ER stress-associated transcription factor XBP1s, based on MS TMT intensity, in thapsigargin (Thap)-treated and control samples, in the NP-40-insoluble, NP-40-soluble, and total protein fraction. adjP: limma adj. P. F. Schematic depicting cellular compartment localization of the different types of condensates.

Multiple proteins were found only in the insoluble pellets but not the soluble supernatent, suggesting they are highly partitioned to insoluble fractions in normal or stressed cell (Scenario 4). These include several proteins involved in RNA splicing, epigenetic regulation, and membraneless organelle assembly.

Phosphopeptides in particular can show prominent re-distributions under cell stress by driving only specific phosphoproteoforms of a protein across solubility boundaries (Scenario 5). As many phosphopeptides in the data are only found within the insoluble pellets, to estimate partition, we considered the stress-induced differential expression of a phosphopeptides normalized by the unmodified protein average in the insoluble fraction (Δ phosphorylation), over the differential partition of the unmodified protein (Δ partition), as an estimate of the phosphorylation-based partition change of a phosphoproteoform in stressed cell (**Figure 3E**). From this analysis, we noticed that the magnitude of stress-induced phosphorylation bias in the insoluble fraction is often greater than the differential partition of the native unmodified protein, consistent with substantial protein regulation by post-translational modification in stressed cells (**Figure 3E**). Second, many proteins show relatively little stress-induced differential partition at the native unmodified protein level (Δpartition^protein^ < threshold) but significant differential abundance of phosphopeptides within the NP-40-insoluble fraction (**Figure 3E**, blue), which contrasts with the behavior of proteins with differential partition but little phosphorylation changes (**Figure 3E**, green). For the larger former group, the data therefore points to the contributions of site-specific phosphorylation status, i.e., phosphoproteoforms, in modulating the function of the insoluble protein pool, or driving stress-induced protein differential partition.

### Phosphoproteome partition under stress involves multiple types of biomolecular condensates

As we identified RBPs including SRRM2 and HNRNPA2B1 with specific phosphorylation-driven partition (**Figure 3E**), consistent with the dynamic assembly of ribonucleoprotein assemblies in stressed cells, we focused on composition of the NP-40-insoluble fraction next and considered the differential abundance of proteins and phosphoproteins within the insoluble fraction after stress.

NP-40-insoluble but SDS-soluble proteins may contain protein aggregates as well as proteins localized to various biomolecular condensates within the cell (**Figure 4F**). We grouped proteins and phosphoproteoforms within the insoluble fractions by their known condensate type annotation in CD-CODE (24) and their differential regulation direction within the insoluble fraction. Notably, proteins and phosphoprotein with significant abundance changes within the insoluble fraction map to diverse types of known nuclear and cytoplasmic biomolecular condensates, including stress granules, P bodies, PML body, nucleolus and nuclear speckles (**Figure 4A**). This implies that the composition of these condensates may vary in prolonged ER stress, and moreover, proteome-wide solubility remodeling affects not only stress granules but various nuclear bodies.

Furthermore, there appears to be a bias in the direction of differential abundance between proteins and phosphopeptides across the condensate types. Whereas total unmodified proteins belonging to most condensate types trend toward decreased abundance within the insoluble fraction, the phosphopeptides largely show bimodal behaviors with both depleted and accumulated phosphoproteoform across the stress granules, P bodies, and an increase in nuclear speckle phosphoproteins under stress (**Figure 4A- B**). This dichotomy is consistent with a complex consequence of the charge-altering effects of phosphorylation that may promote electrostatic attraction or repulsion, with phosphorylation on some proteins driving accumulation toward the condensates and others driving depletion from condensates. The nuclear speckle presents an exception in our observations, with most phosphoproteins of proteins annotated to be in the nuclear speckle showing increased abundance within the insoluble fraction.

To further investigate the different solubility behaviors of unmodified proteins and phosphoproteins, we performed Gene Ontology overrepresentation analysis (ORA) on the proteins and phosphoproteins that show significant differential expression in the insoluble fraction after stress but not in the soluble fraction, indicative of insoluble compartments specific behavior. Interestingly, the analysis reveals drastically different set of gene names between the unmodified and phosphorylated protein analysis, with unmodified proteins largely enriched in the ER lumen, extracellular matrix, and ribosome, which are contributed by proteins not known to be part of biomolecular condensates, including thrombospondin, fibrillin, among others. Therefore, the differential solubility behavior of unmodified proteins is largely driven by abundant ER, secretory, and fibrillar proteins, indicating these proteins may become NP-40- insoluble because of aggregate formation due to secretory pathway protein misfolding (**Figure 4D**, top). On the other hand, the phosphoproteins with significant insoluble fraction abundance under stress are enriched in nuclear speckle, chromosome, and spliceosome (**Figure 4D**, bottom). Therefore, proteome solubility changes under prolonged ER stress show both phosphorylation-independent and phosphorylation-dependent behaviors reflecting distinct biophysical events, with phosphorylation-independent changes likely representing assembly while phosphorylation-dependent solubility behaviors associated with dynamic biomolecular condensate remodeling.

To investigate further, we performed a network analysis of phosphopeptides and their proteins within the insoluble fraction, mapped to STRING database protein interactions (25). The main network shows a clustering of protein neighborhoods based on protein-protein interactions, which also show signs of concordant directions in average post-stress phosphopeptide changes within the insoluble fraction (**Figure 4C**). For instance, module 1 connects multiple known protein-protein interactors that are annotated to reside in the nucleolus, including RPS7 and SERBP1, and SEC61B, which show downward trends in stressed cells. Conversely, module 4 contains multiple interaction partners in the nuclear speckles, including SRSF7, SRRM1, SRRM2, CCNL1, and U2SURP, that are up-regulated in thapsigargin treatment. This suggests protein phosphorylation within insoluble proteome may be coordinately regulated based on protein interactions or localization.

### Phosphorylation-driven condensate behaviors of splicing related proteins in stressed cells

We next considered individual splicing related phosphoproteins with significant stress-induced phosphopeptide abundance within insoluble fractions. These splice factors and splicing-related proteins span different annotated categories including core spliceosome components, hnRNPs, and SR proteins. Among these proteins, multiple SR proteins including SRSF6, SRSF10, SRSF3, SRSF4, and SRSF7 show increases in the insoluble fraction, consistent with a stress-induced dynamic shuttling from soluble compartments into condensates. On the other hand, several hnRNP proteins and spliceosomal helicases including DDX6, DHX16, HNRNPA2B1, and HNRNPUL1 show decreased phosphopeptides abundance within insoluble fractions under stress, consistent with these proteins shuttling away from condensates in stressed cells (**Figure 5B**). The contrasting regulations of SR and hnRNP families of splice factors is notable, as the differential sequence properties of SR and hnRNP proteins play a role in their differential compartmentalization: whereas SR proteins are enriched in nuclear speckles, hnRNPs are either unenriched or depleted from nuclear speckles and often partition with paraspeckles (26). Their contrasting behaviors in the NP-40-insoluble fraction indicate that ER stress exerts a differential effect on the organization of splice factors across nuclear bodies.

**Figure 5:**
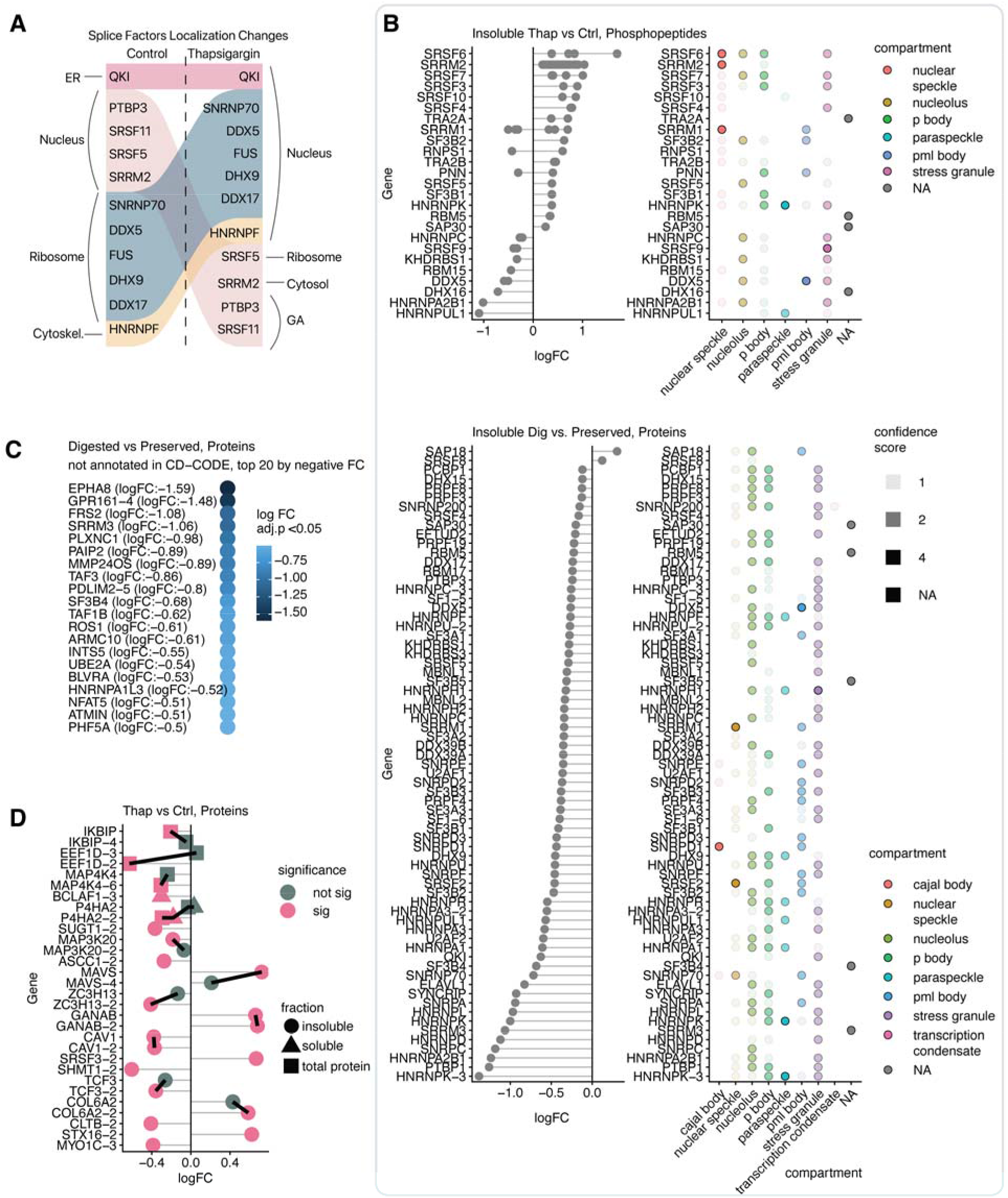
Phosphorylation-dependent partition of splice factors and splicing-related proteins. A. Results from the analysis of a subcellular proteomics experiments highlighting the differential localization of several splice factors under thapsigargin (BANDLE differential localization probability > 0.99). The labeled subcellular compartment refers to classification based on known compartment markers, which do not include known condensates. Attention should be given to broad redistribution from or to the nuclear compartment. B. Top: Changes in protein phosphorylation across known splice factors (limma adj. P < 0.05; thapsigargin/control) in the insoluble fraction. Each data point represents a phosphopeptide. Annotations to the right show the CD-CODE annotated condensate membership of each protein. Confidence score denotes confidence assignment from CD-CODE. Bottom: Differential abundance of proteins in the NP-40 insoluble fraction after RNase treatment (RNA-digested (Dig.) vs. RNA-preserved). The majority of RBP shows negative abundance in the insoluble fraction after RNA digestion, indicating their solubility behavior reflects ribonucleoprotein granule formation. Confidence score denotes confidence assignment from CD- CODE. C. The differential abundance of additional proteins in the NP-40 insoluble fraction after RNase treatment. The top 20 proteins with the most negative logFC after RNase and which are not currently annotated as a condensate member in CD-CODE are shown, including sevearl splice factors, SRRM3, and PDLIM2. D. Differential abundance (thapsigargin (Thap)/control (Ctrl) of a number of protein isoform pairs in the insoluble fraction. Protein isoforms are shown if any of the isoforms from the same protein is significantly altered (limma adj. P < 0.05)

Interestingly, for a majority splicing-related proteins, multiple phosphoproteoforms of the same protein share largely coherent behaviors (**Figure 5B**), suggesting these phosphorylation sites may function via tuning protein or IDR net charges (9). However, proteins where different phosphorylation sites show contrasting stress response also exist (RNPS1, PNN, SRRM1) which may indicate site-specific functions. On the other hand, the number of significant phosphorylation changes with the native unmodified splicing-related proteins, where only one protein (SRSF3-2) passes the significance threshold when comparing stress-induced abundance change within the insoluble fraction. This is consistent with the analysis on overall foldchange distributions of proteins annotated to be in nuclear speckles (**Figure 4A**) and corroborates that phosphorylation profiling is critical in detecting dynamic splice factor partition changes.

To corroborate that the solubility behavior of these splicing-related RBPs represent bona fide membership in biomolecular condensates within nuclear bodies, we treated cell lysate with RNase prior to NP-40 extraction and compared RBP abundance in the NP-40-insoluble fraction following RNA digestion. RNA is required for the assembly of a number of cytoplasmic and nuclear bodies (27,28). Consistent with this, we find that the majority of RBPs show decreased residency in the NP-40-insoluble fraction (i.e., become more soluble) when RNA is digested (**Figure 5C**), thus strongly supporting that the insolubility of RBPs reflect their membership within ribonucleoprotein assemblies. In parallel, we also identify multiple proteins with significantly increased solubility (RNase digested vs. control logFC < –0.3, limma adj. P < 0.05) upon RNase treatment that are not currently documented within CD-CODE (24), but featuring high predicted condensate membership likelihood by PICNIC (29), including TAF3, HNRNPA1L3, and SRRM3, nominating these proteins to be uncharacterized stress-responsive members of membraneless organelles in the cell (**Figure 5C**)

We note that among the exceptions to this RNA-dependent behavior of splicing-related RBPs are the phosphopeptides of SRRM2, a protein that acts as a scaffold for nuclear speckles (6,30), and which became more insoluble in the proteomics experiments when RNA is digested (**Supplementary Figure S2)**. Prior work has shown that SRRM2 forms a few larger nucleoplasmic aggregates when nuclear speckles are dissolved by RNA removal (27), consistent with its baseline detergent solubility being tied to RNP granule membership. Therefore, these results support that the observed solubility change represent bona fide ribonucleoprotein granule behavior.

### Differential condensate partition of nuclear speckle components in prolonged ER stress

Our data and analysis center on the nuclear speckle and associated splice factors as being a nexus of stress-induced remodeling, with multiple SR proteins showing significant alterations in their solubility behavior upon stress. To investigate their stress response by orthogonal techniques, we performed immunofluorescence imaging of SRSF1, SRSF2, SRSF3, SRSF4, SRSF5, SRSF6, SRSF10, and SRSF11 in normal and stressed cells. Nuclear speckles, for which SRSF1 and SRSF2 are commonly used as canonical markers, can be seen as distinct puncta within the nucleus of AC16 cells and quantified as the proportional intensity of dense area in the images (**Supplementary Figure S3**). The imaging analysis shows a significant increase in relative puncta partition for SRSF5 and SRSF6 upon stress suggesting preferential movements toward condensate bodies. On the other hand, SRSF4 shows significantly lower dense area to total area ratios in fluorescence intensity upon stress, and a significant cytosolic fluorescence can be observed, suggesting shuttling out of the nucleus (**Supplementary Figure S3**).

In parallel, we visualized the nuclear speckle scaffold protein SRRM2 in stressed cells. Within the solubility proteomics data, SRRM2 stands out the protein with the most discovered phosphopeptides (**Figure 1D**), whereas all quantified SRRM2 phosphopeptide show significant increases after stress within the insoluble fraction, indicating preferential association with condensate (**Figure 5A**). As opposed to most RBPs, the removal of RNA by RNase treatment prior to NP-40 extraction renders SRRM2 phosphopeptides less soluble, indicating RNA-bound SRRM2 is more soluble and that under stress response, phospho-SRRM2 are excluded from RNA. To test whether the insolubility of SRRM2 upon stress reflects aggregate formation, we performed immunofluorescence imaging co-localization analysis between SRRM2 and SRSF2 in normal and stressed cells (**Supplementary Figure S4**). In normal cells, SRRM2 signals show strong co-localization with SRSF2 and DAPI, indicating a virtually exclusive nuclear speckle localization. Upon ER stress, this co-localization weakens and SRRM2 distribution becomes more diffuse, but we did not observe the appearance of large SRRM2 aggregates that would suggest nuclear speckle dissolution (27). On the contrary, under expression of G3BP1-SNAP, we find evidence of co-localization between SRRM2 and G3BP1 in discrete puncta, suggesting SRRM2 translocates out of the nucleus and may associate with other biomolecular condensates (**Supplementary Figure S4**). Prior work in the literature has suggested that when certain RBPs mislocalize to the cytoplasm and encounter a lower RNA concentration, they may form solid-like condensates (33).

To further evaluate the evidence of translocation of SRRM2, we re-analyzed of a prior subcellular spatial proteomics dataset of protein subcellular localization under 16 hours of thapsigargin treatment to assess RBP localization (**Figure 5A**). The data reveals a movement of SRRM2 from the nucleus to the cytosol at 16 hours of ER stress, corroborating SRRM2 may be exported from the nucleus under stress. Interestingly, the data also reveals a significant differential localization of two SR proteins (SRSF11 and SRSF5) away from the nucleus fraction toward the ribosome, cytosol, and Golgi apparatus fractions; whereas the several RNA helicases including DDX5, DHX9, and DDX17 show movement from the ribosome fraction toward the nucleus. As the ribosome and Golgi fractions were classified based on their co-sedimentation with canonical fraction markers and contained condensates with similar sedimentation coefficients as these organelles, the spatial proteomics results are consistent with a localization of SR proteins toward condensates and RNA helicases away from condensates under prolonged ER stress.

Finally, our experiments also detected a number of protein isoforms with differential (|logFC| > 0) abundance in total, soluble, and insoluble proteome fractions under stress, which may reflect the outcome of altered splicing programs (**Figure 5D**).

## Discussion

Here we report a comparative solubility proteomics study using a new hybrid bottom-up and chemical middle-down proteomics strategy, analyzing the behavior of of 8,740 proteins and 31,647 phosphopeptides in normal cells and cells under prolonged ER stress. To increase protein coverage especially at lysine-rich IDR, we implemented a chemical dimethylation method to extend tryptic peptides in the proteomics and phosphoproteomics experiments. While on paper this method achieves the same enzyme cutting rule of Arg-C, our method takes advantage of the low cost, robustness, and actual specificity and selectivity of trypsin over alternative proteases. In our data, middle-down peptides led to increased coverage of phosphorylation sites, including along disordered and low complexity regions of multiple splicing-related proteins (**Figure 6**). We envision that the strategic use of lysine and arginine blockade, as well as the continued development of long peptide detection via middle-down and top-down proteomics, will provide valuable tools for resolving proteoforms in IDR, and help distinguish the site specific and charge-tuning effects of phosphorylation events in condensate partition.

**Figure 6:**
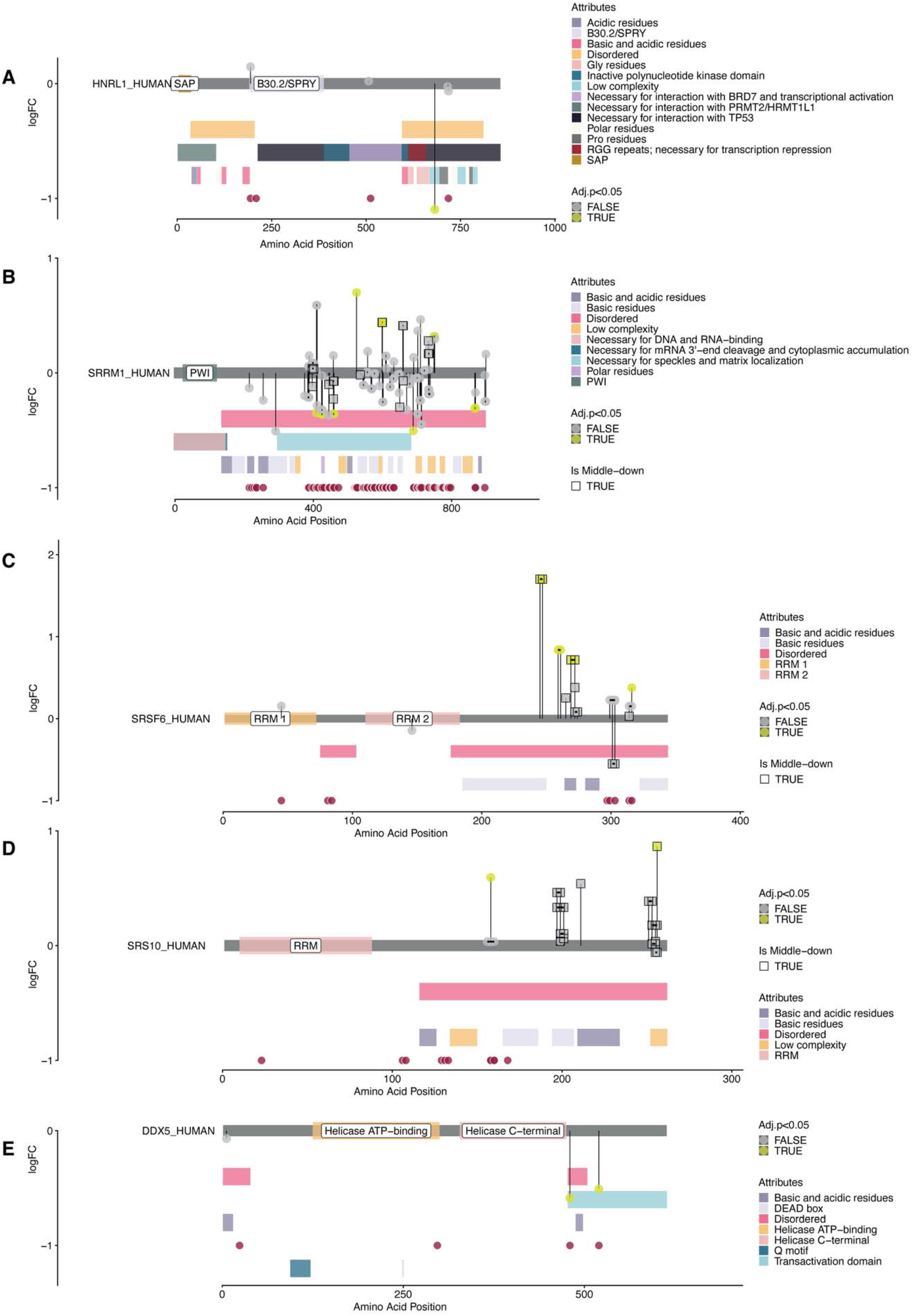
Phosphopeptides of splicing-associated proteins show concerted stress-induced changes within NP-40 insoluble fraction. A. Protein track of HNRNPUL1, denoting the differential abundance in logFC (thapsigargin/control; y-axis) and position (x-axis) of quantified phosphopeptides within the NP-40-insoluble fractions. Square data points denote phosphopeptides identified only via middle-down experiments. Yellow data points: limma adj. P < 0.05. Annotation track positions and colors denote protein domains, motifs, and sequence features on UniProt. Link between data points connect multiple phosphorylation sites within the same identified phosphopeptide. B. As in A, but for SRRM1. C. As in A, but for SRSF6. D. As in A, but for SRSF10. E. As in A, but for DDX5.

A major goal of proteomics analysis is to survey the inventory and functional states of the proteome of the cell in order to find disease correlates. In recent years it has become increasingly clear that protein abundance only represents one dimension of such changes and that protein function is also regulated by other dynamic parameters including localization and modifications. Mapping the changes in protein solubility has been frequently done under neurodegenerative disease and aging using various detergents, including NP-40, sarkosyl, and SDS (34,35). Our study is distinguished from prior works that largely focused on protein aggregates and amyloid fibril behaviors, and builds on recent solubility proteomics advances (22) demonstrating NP-40 solubility, especially in conjunction with RNase sensitivity analysis, provides a systematic means to survey proteome-wide biomolecular condensate membership, and moreover extends this approach toward probing the stress-induced dynamics of global protein solubility using deep middle-down phosphoproteomics profiling.

Compared to membrane-bound organelles and stoichiometric protein complexes, biomolecular condensates can assemble and disassemble dynamically, making it a particularly suitable mode of molecular compartmentalization in the context of cellular regulation and stress response (36). The stress-induced partition of proteins in and out of condensates may be driven by protein concentration, charge modification such as by phosphorylation or arginine methylation, or the cellular environment. For instance, whereas low concentration of RNA may support or indeed be required for the formation of condensate, RNA concentration above a certain threshold may suppress protein condensate formation due to charge repulsion (37,38). Prior proteomics works have focused on the context-specific changes in protein membership within stress granules and under acute (30 min to 1 hr) stress and integrated stress response (ISR) induction, but noted relatively minor difference in the pre- vs. post- stress G3BP1 interactome (39,40). As it is known that cellular proteomes continue to evolve past initial stress induction and ISR signaling and time-specific changes in ER stress may become most prominently observed at the protein level only under prolonged stress at up to 16-24 hours (41), we investigated a late phase stress response to capture the proteome-wide effect of stress on protein solubility as cells adapt to restore proteostasis. A striking observation is that the NP-40-insoluble proteome fraction contains over 7,000 proteins, and moreover up to 241 proteins change significantly in their partition coefficients under ER stress. Our results therefore indicate a proteome-wide rearrangement of solubility, including the association and disassociations of RBP and splicing-related proteins to or from biomolecular condensates, as a prominent feature of the stressed proteome. Moreover, stress-induced protein level measurements are dramatically different between the insoluble and the soluble proteomes, indicating protein partition dynamics is at least partially orthogonal from gene expression changes in ER stress, and represents an under-investigated dimension of proteome regulation.

Nuclear speckles are dynamic ribonucleoprotein assemblies within the nucleus that play critical roles in modulating the availability and function of splice factors and other RBPs (14,42). Nuclear speckles integrate a variety of cellular cues to orchestrate splicing output (42,43) and have in recent years been implicated in stress response. For instance, stress response genes are enriched among RNAs that engage efficiently with speckles-resident splice factors (43), nuclear speckles may act as a sponge to sequester transcripts during global transcription inhibition to prevent translation, analogous to the role of stress granules in the cytoplasm (44), whereas speckles may also act in the contexts of ER stress through an XBP1s-SON axis to amplify the expression of proteostatic genes (45). Protein phosphorylation has emerged as a central determinant of nuclear speckle organization and function (14,42). For instance, phosphorylation drives the nucleocytoplasmic shuttling and nuclear speckle targeting of SRSF1 (46,47); whereas hypoxia but not heat shock leads to a hyperphosphorylation but overall down-regulation of SRSF6 while promoting nuclear speckle dissolution (18). Our proteome-wide investigation shows that that multiple SR and hnRNP proteins and nuclear speckle components show accumulation within the condensate-rich proteome fractions in prolonged ER stress. The observation of increased insoluble fraction accumulation of phosphopeptides of SR proteins is consistent with greater speckle localization of these proteins in stressed cells, whereas imaging also supports an increased nuclear body residency of SRSF5 and SRSF6.

The phosphorylation of SR proteins may reflect a bimodal response between acute and prolonged stress. For instance, a recent work found that acute heat shock and arsenite (1 hr) both decreased overall SR protein phosphorylation together with mRNA retention and translation inhibition (17). On the other hand, prolonged hypoxia (24 hours) leads to SR hyperphosphorylation (18). Our data here in turn shows that prolonged ER stress and UPR led to the accumulation of SR protein and SRRM2 phosphoproteoforms within insoluble fractions. This hints at a phased nuclear speckle response where acute stress response leads nuclear speckle dephosphorylation and translation inhibition, but prolonged stress response initiates distinct modality of remodeling.

SRRM2 is one of the handful RBP and splicing-related proteins showing an exceptional behavior, where the removal of RNA by RNase treatment prior to NP-40 extraction renders SRRM2 less soluble. The accumulation of phospho-SRRM2 under prolonged ER stress indicates they are excluded from RNA. This is consistent with prior work showing overexpression of DYRK3 to phosphorylate nuclear speckle components leads to less polyA+ mRNA retained in the nucleus (17), and suggests that the global phosphorylation state of SRRM2 may act as a switch to control nuclear speckle RNA engagement during stress response. Taken together, these results reinforce the notion that splicing machineries actively participate in and constitute a central part of cellular stress response (14).

### Limitations of the study

Our results further highlight that solubility partition behaviors are highly linked to members of nuclear bodies. However, as the NP-40-insoluble fraction is enriched in multiple types of biomolecular condensates and aggregates, the described method does not distinguish the membraneless organelle locations of proteins and their permutations under stress. For instance, paraspeckles have significant crosstalk with nuclear speckles and may accumulate under stress (31), whereas also nuclear stress bodies may form that phosphorylates SRSFs (48). Many condensate-forming proteins also participate in multiple condensates (24,36). The interactions between membraneless compartments may not result in net difference in protein solubility behavior. Moreover, membraneless organelles contain sub-phases (42,49–51), how distribution of phases are reflected in protein solubility differences is not addressed.

Despite these limitations, the results show that proteome-wide solubility partition is a prominent feature of proteome stress response. Applied to other system, this method may inform on whether an up-regulation of a total protein pool occurs in conjunction with the protein residing in its normal functional compartment.

## Methods

### Cell culture and ER stress induction

AC16 human cardiomyocyte cells (Millipore) were cultured in DMEM/F12 supplemented with 10% fetal bovine serum (FBS) without antibiotics and maintained at 37 °C in a humidified incubator with 5% CO . Cells used were at passage numbers ≤10 and grown for 3 days prior to treatment. Endoplasmic reticulum (ER) stress was induced by treating cells with 1 µM thapsigargin for 16 h. Cells were harvested by treatment with 0.25% trypsin, pelleted by centrifugation at 300 × g for 3 min, washed three times with phosphate-buffered saline (PBS). The cell pellets were immediately snap-frozen and stored in liquid N_2_.

### Solubility fractionation of AC16 cell lysates

Lysis buffer was prepared in phosphate-buffered saline (PBS) with 1 U/ml RNase inhibitor (RNasin plus), cOmplete protease inhibitor, phosphotase inhibitor (PhosSTOP), 1.5 mM MgCl_2_, 2 mM NaF, 2 mM Na_3_VO_4_, 2 mM Na_4_P_2_O_7_ and 10 mM CH_3_CH_2_CH_2_COONa. Frozen AC16 pellets were thawed on ice for 10 minutes and 200 µl of lysis buffer was added to the AC16 cell pellets derived from approximately 5 × 10 cells. The cell suspension was mechanically disrupted by three freeze-thaw cycles using liquid N_2_. For each biological replicate (control and thapsigargin treated), approximately 22–25 × 10 cells were used. Protein concentrations were determined by the Rapid Gold BCA assay. The lysates were normalized to 3.5 mg/ml with lysis buffer and split into three aliquots: (1) total proteome lysate, (2) RNA-preserved lysate and (3) RNA-digested lysate. The aliquots for total proteome lysate and RNA-digested lysate were treated with Benzonase (1:100 v/v) and RNase cocktail (1:50 v/v) respectively. All three aliquots were incubated in a cold-room at 4 °C with a shaking platform at 500 rpm for 30 minutes. The aliquots for RNA-preserved and RNA-digested lysates were added with NP-40 to a final concentration of 0.8%. The insoluble pellets were isolated with ultracentrifugation at 100,000g for 25 minutes at 4 °C and the supernatants were collected as soluble fractions. The insoluble pellets were washed gently three times with lysis buffer containing 0.8% NP-40 and then resuspended in lysis buffer containing 1% SDS and 0.25 U/ml of Benzonase. The aliquot for total proteome lysate was added with SDS to a final concentration of 1%. The insoluble fractions and total protein fractions were incubated at room temperature for 15 minutes and followed by heat treatment at 90 °C for 5 minutes.

### Protein Digestion

All samples were diluted with an equal volume of sonication buffer consists of 1% w/v sodium deoxycholate, 5 mM tris(2-carboxyethyl)phosphine (TCEP), 30 mM chloroacetamide (CAA), 1 mM MgCl_2_, 10 U/μl Benzonase in 100 mM HEPES, and sonicated in a Bioruptor for 15 cycles (30 s on and 30 s off) at 4 °C. The protein concentrations were quantified using the Rapid Gold BCA assay. Proteins were reduced and alkylated with 1.7 mM TCEP and 5 mM CAA for 30 minutes at 37 °C and split into two equal aliquots for non-dimethylation and dimethylation.

Samples not intended for demethylation were subjected directly to SP3 trypsin/LysC digestions. Briefly, Sera-Mag SpeedBeads (10:1 w/w beads to protein ratio) were added to the protein samples and immediately followed by the addition of 100% ethanol to bring the final concentration of ethanol to 70% v/v. The mixtures were vortexed for 20 minutes at room temperature (VWR mixer at speed 7). After magnetic separation, the supernatant was discarded and the beads were washed with 70% ethanol and resuspended in 200 µl of 100 mM HEPES buffer (pH 8.0) and incubated with trypsin/LysC (1:50 enzyme to protein ratio w/w) at 37 °C overnight.

For samples to be demethylated, Sera-Mag SpeedBeads (10:1 w/w beads to protein ratio) were added to the proteins, followed by the additional of 100% ethanol to bring the final concentration of ethanol to 80% v/v. The mixtures were vortexed for 20 minutes at room temperature to allow protein binding to the beads. After magnetic separation, the supernatant was discarded and the beads were washed with 80% ethanol for 3 times. The beads were resuspended in freshly prepared dimethylation buffer (30 mM formaldehyde, 15 mM sodium cyanoborohydride and 100 mM HEPES, pH 8.0). The mixture was incubated at 37 °C for 1 hour in a Thermo Scientific ThermoMixer with shaking, followed by another hour of incubation with an additional aliquot of 3 µl formaldehyde (2 M) and 3 µl sodium cyanoborohydride (1 M) at 37 °C. The reaction was quenched by 2 µl of 50% (v/v) hydroxylamine for 10 minutes (at room temperature). Additional sera-Mag SpeedBeads was added (5:1 w/w beads to protein ratio) to the mixture followed by an addition of100% ethanol to a final concentration of 80% (v/v) ethanol. The mixture was vortexed at room temperature for 15 minutes. The supernatant was discarded after magnetic separation and the beads were washed with 80% ethanol for 3 times. The beads were resuspended in 200 µl of 100 mM HEPES buffer (pH 8.0) and incubated with trypsin/LysC (1:50 enzyme to protein ratio w/w) at 37 °C overnight.

All proteolyzed peptides were dried with SpeedVac and desalted with Pierce desalting spin columns.

### TMT-labeling

For the soluble (RNA-Preserved and RNA-Digested) and total proteome samples, non-dimethylated and dimethylated samples were labeled (TMTpro16-plex) in separate groups. There were a total of 30 samples and they were randomly divided into these two TMT-labeling groups (non-dimethylated and dimethylated). Briefly, after digestion, the desalted peptides were reconstituted in MS-grade water and concentrations were determined using a NanoDrop spectrophotometer. Peptides (100 µg) of each biological replicates were labeled with TMTpro 16-plex reagents in 100 mM HEPES and 20% acetonitrile (v/v) at a 1:2 (peptide:TMT) w/w ratio for 1 h at room temperature. The reaction was quenched with 2 µl of 5% hydroxylamine at room temperature for 15 minutes. The samples were then acidified with trifluoracetic acid (TFA, final 2%). The labeled peptides within the same TMT-labeling group were pooled and dried with a SpeedVac.

The above TMT-labeling procedures were similarly applied to the labeling of insoluble proteins (non- dimethylated and dimethylated).

The pooled TMT-labeled peptides from each TMT-labeling group were reconstituted in 100 mM HEPES. One-tenth of the peptides were subjected directly to high-pH fractionation. The remaining peptides were used in phosphopeptide enrichment.

### High pH reversed-phase fractionation

The non-phospho-enriched peptides were fractionated with Thermo High-pH fractionation kit into eight fractions and dried with a SpeedVac.

### Phosphopeptide enrichment

The pooled TMT labeled peptides were desalted with Pierce Peptide Desalting Spin column and dried using a SpeedVac. Phosphopeptides in each pooled sample were first enriched with Titansphere Phos-TiO Tip (GL Sciences) according to the manufacturer’s protocols. The enriched phosphopeptides were sequentially eluted from the Phos-TiO Tip with 5% ammonium hydroxide and 5% pyrrolidine. The collected phosphopeptides were dried with a SpeedVac and desalted with Pierce C-18 spin columns. The second phosphopeptide enrichment was performed on the flow through collected from the first Phos-TiO Tip using the High-Select™ Fe-NTA Phosphopeptide Enrichment Kit (Thermo Fisher Scientific) following the manufacturer’s instructions. The collected phosphopeptides were dried with a SpeedVac and desalted with Pierce C-18 spin columns.

### Mass spectrometry

#### Non-phosphoenriched samples

TMTpro-16 plex labeled high-pH fractionated peptides were resuspended in solvent A (0.1% FA in 100% LCMS-grade water) and injected on to an EasySpray^TM^ C18 analytical column (PepMap C18, 3 μm, 100Å, 75 μm x 15 cm). The peptides were separated by Vanquish Neo UHPLC system (Thermo Fisher Scientific) using solvent A (100% LC-MS water with 0.1% FA) and solvent B (80% LC-MS acetonitrile with 0.1% formic acid). Peptides were separated by 90 minutes gradient: solvent B from 2-67% for 66 minutes, 24-55% for 16 minutes, 55-99% for 1 minute, and held at 99% for 5 minutes. Mass spectrometric data were acquired in data-dependent mode on the Orbitrap Exploris 480 (Thermo Fisher Scientific) operated in positive ion mode. The spray voltage was operated in static mode and the capillary temperature 275 °C. The full MS1 scan were acquired in profile mode with a mass range of 375-1600 m/z (375-1800 m/z for dimethylated samples) at a resolution of 120,000 using one micro scan, auto maximum injection time, an automatic gain control (AGC) target of 300% and RF lens of 50%. The precursor intensity threshold as 3000 and charge states selected as 2-5 (2-8 for dimethylated samples). MS2 scans were acquired at the orbitrap resolution of 30,000 using an isolation window 0.7 with auto maximum injection time and HCD with normalized collision energies at 36. TurboTMT was enabled for TMTpro reagent quantification with a normalized AGC target as 200%.

#### Phosphoenriched samples

TMT labeled TiO_2_ and Fe^3+^-NTA enriched phosphopeptides were resuspended in solvent A and injected onto an EasySpray^TM^ C18 analytical column. Peptides were separated using Vanquish Neo UHPLC system (Thermo Fisher Scientific) using solvent A and solvent B as described above. Phosphopeptides were eluted by 120 minutes gradient: solvent B was held isocratic at 2% for 1.1 minutes, with stepwise increase from 2-24% for 96 minutes, 24-99% for 16 minutes, 55- 99% for 1 minute, and held at 99% for 5.9 minutes. Mass spectrometric data were acquired in data-dependent mode on the Orbitrap Exploris 480 (Thermo Fisher Scientific) mass spectrometer using the same acquisition parameters as described above in non-phosphoenriched samples.

#### Cell transfection and treatment

Plasmids encoding G3BP1 with serine at position 149 (G3BP1_S149; control), the phosphomimetic G3BP1_S149E and the phospho-mutant G3BP1 S149A were purchased from GenScript. All plasmids are fused to SNAP-tag on C-terminus. AC16 cells were cultured DMEM/F12 supplemented with 10% fetal bovine serum (FBS) without antibiotics and maintained at 37 °C in a humidified incubator with 5% CO_2_. At 60% confluency, cells were transfected with the respective plasmids using Lipofectamine 2000 as per the manufacturer’s instructions. Twenty-four hours after medium was changed, transfected cells were treated with 1 μM thapsigargin for ER stress and with DMSO for control. Control and treated cells were incubated for 16 h at 37 °C in a humidified incubator with 5% CO . Cells were then washed three times with PBS and incubated with SNAP-tag substrate (SNAP-Cell 647-SiR) for 30 minutes in the incubator at 37 °C in 5% CO_2,_ according to the manufacturer’s protocol.

#### Immunofluorescence and confocal microscopy

Cells were fixed with 4% formaldehyde in PBS for 10 minutes at room temperature and washed three times with PBS. Cells were permeabilized with 0.1% triton-X100 in PBS for 10 minutes and then blocked with 3% BSA in PBS for 2 h at room temperature. Cells were incubated overnight with primary antibodies (SRRM2, SRSF1, SRSF2, SRSF3, SRSF4, SRSF5, SRSF6, SRSF10, SRSF11) prepared in blocking buffer at 4 °C overnight. After three times with PBS, cells were then incubated with secondary antibodies (Host specific Alexa Flour 488/594) for 1hour at room temperature. Cells were washed three times with PBS and incubated with DAPI solution for 5 minutes. For imaging, cells were mounted using ProLong Gold Antifade mountant. Cell images were captured by EVOS™ M5000 Imaging System with 20X objective (**Supplementary Figure S1**) and 3I Marianas Spinning Disk Confocal with 63X oil objective using SlideBook software (**Supplementary Figure S3** and **Supplementary Figure S4**). Cell images were further processed with Fiji.

#### Database search, analysis, statistics

TMT data were processed using FragPipe. Peptides were searched against UniProt human database (UP000005640 reviewed with isoforms, downloaded on 2025-09-05).

#### Non-phosphoenriched, Non-dimethylated Samples

Variable modifications were set to M 15.9949 (max = 3), N-terminal 42.0106 (max =1), and N-terminal 304.20715 (max=1). Fix modifications were set to C 57.02146 and K 304.20715. Enzyme was set to “stricttrypsin” with 2 missed cleavages allowed. Peptide length: 7-50; peptide mass range: 200 to 5,000. Max variable mods on a peptide: 3, max combinations: 5,000. <u>Non-phosphoenriched Dimethylated Samples</u>: same as non-dimethylated samples except enzyme specificity was set to “argc” with 1 missed cleavage. Max variable mods on a peptide: 5, max combinations: 10,000. Fixed mods include K 28.0313 and C 47.02146. <u>Phosphoenriched, Non-</u> <u>dimethylated Samples</u>: same as non-phospho-enriched samples with the following modifications: max variable mods on a peptide: 5, max combinations: 10,000, STY 79.96633 (max = 3). <u>Phosphoenriched,</u> <u>Dimethylated Samples</u>: Same as non-phospho-enriched demethylated samples with the following modifications: STY 79.96633 (max = 3).

Statistical analyses for differential expression for protein and phosphopeptides were performed with R limma package with multiple testing correction. GO enrichment was performed with R clusterProfiler.

## Acknowledgement

BP was supported by AHA Postdoctoral Fellowship 25POST1372064. M.P.L. was supported by National Institutes of Health (NIH) grants R01-HL169473, R01-HL141278, and R01-GM144456. E.L. was supported by NIH grant R35-GM146815 and R01-HL169473.

---

**Supplementary Figure S1:**
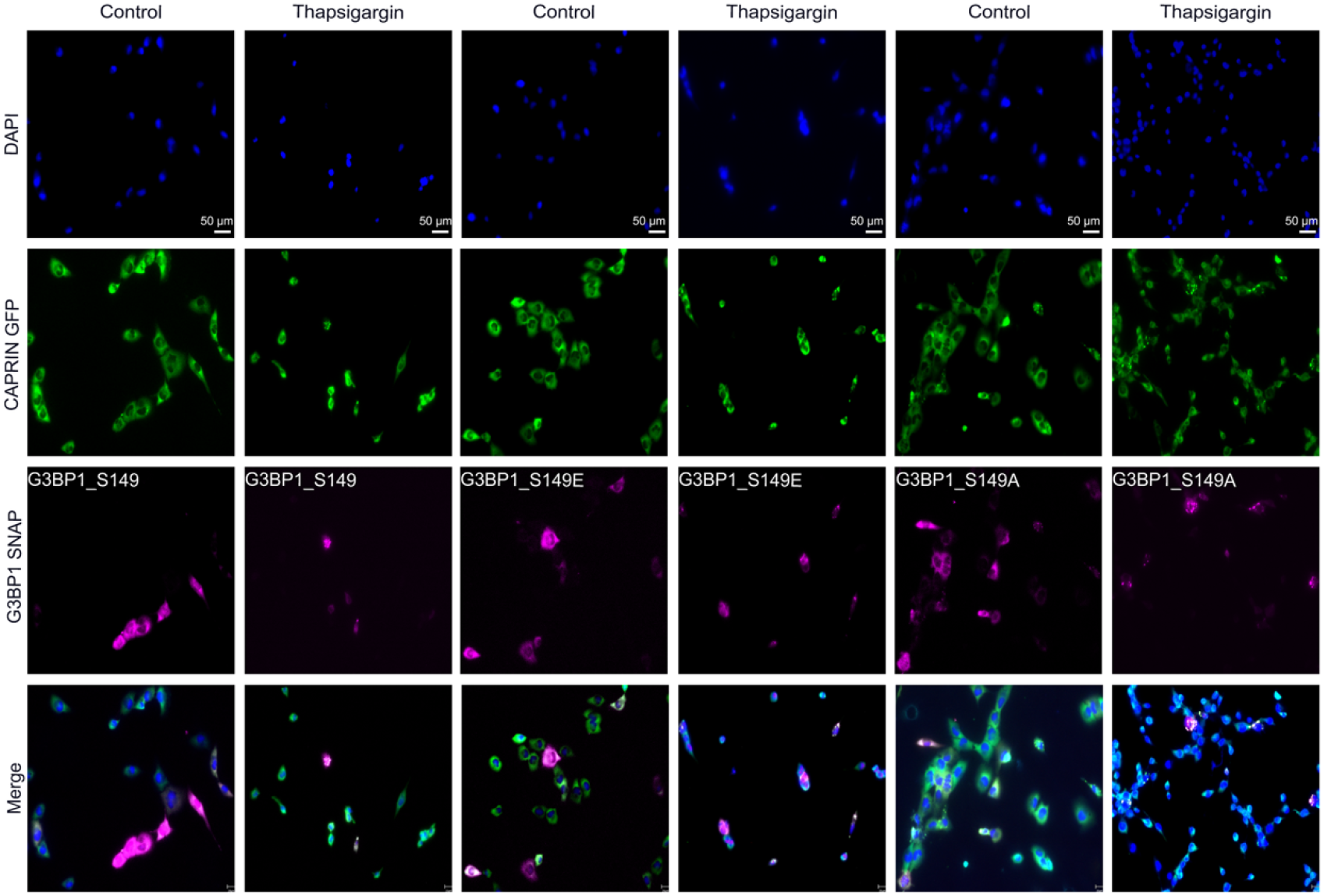
Stress granule formations under prolonged ER stress model. Microscopy images showing puncta formation in control and thapsigargin-treated AC16 cells. AC16 cells are transfected with G3BP1, G3BP1-S149E, and G3BP1-S149 fused to SNAP tags, and either untreated or treated with 1 µM thapsigargin for 16 hours. Blue: DAPI; green: Caprin-1; magenta: SNAP 647-SiR. Scale bar: 50 µm.

**Supplementary Figure S2:**
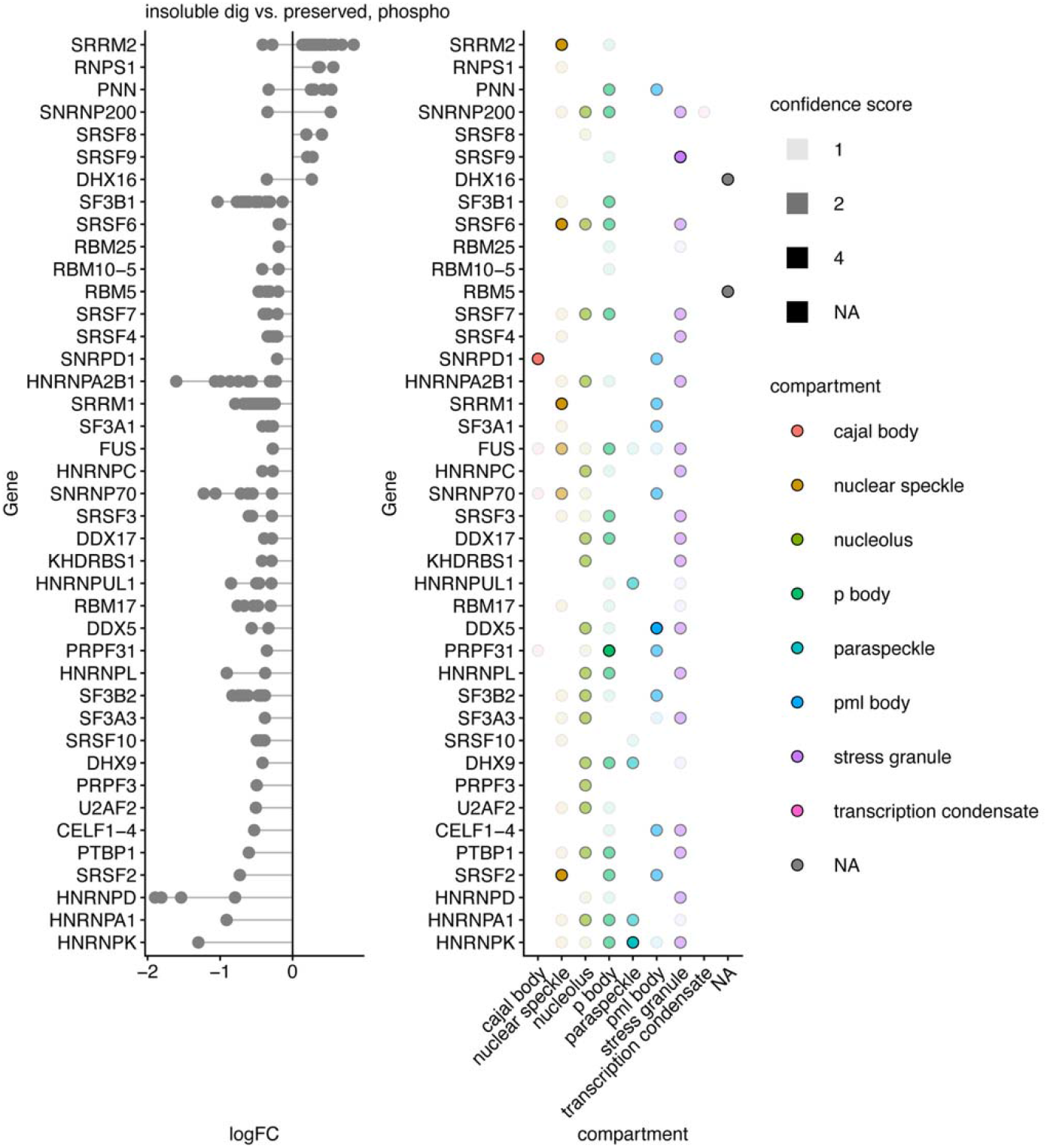
RNase sensitivity of the solubility behaviors of splice factor phosphopeptides. Dot plots showing the logFC within the insoluble pellets of selected splicing-related phosphoproteins with or without RNase treatment. Bona fide ribonucleoprotein granules are expected to be RNase sensitive, i.e., become less enriched in the NP-40-insoluble fraction following RNase treatment. x-axis: logFC RNase/Control. Each data point represents a phosphopeptide. Map on the right represents the CD-CODE biomolecular condensate membership annotation of each protein.

**Supplementary Figure S3:**
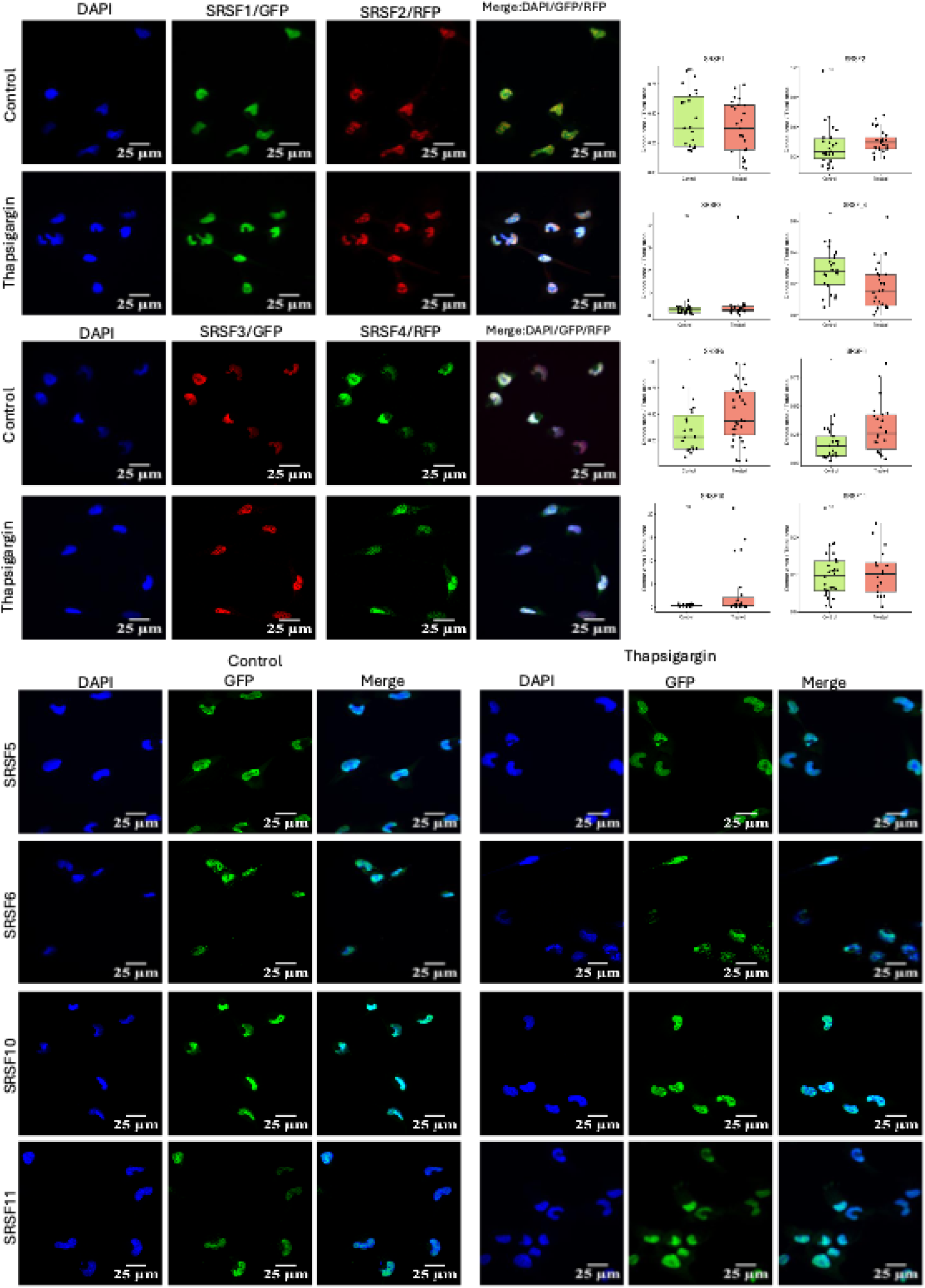
Differential condensate formation of SRSF proteins under stress. Fluorescence microscopy images showing the condensate formation of SRSF family splice factors in normal and stressed AC16 cells. Blue: DAPI; green: GFP; red: RFP. Quantification of puncta is performed using dense area over total area in the images. Scale bar: 25 µm.

**Supplementary Figure S4:**
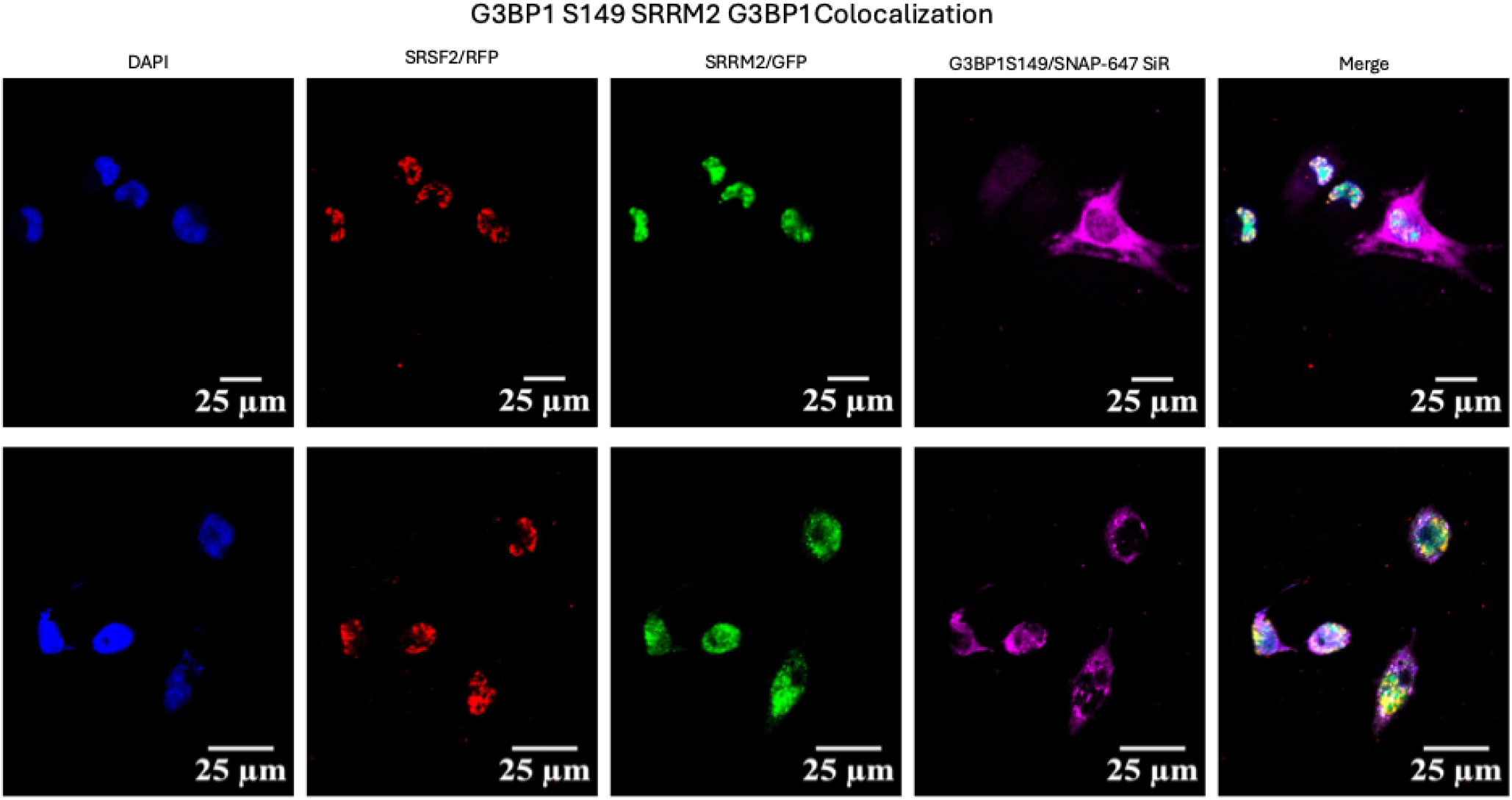
Nuclear export of SRRM2 in stressed cells. Fluorescence microscopy images showing the diffuse and extra-nuclear signals of SRRM2 (green) upon thapsigargin treatment. Blue: DAPI; red: SRSF2 (nuclear speckles marker); purple: G3BP1 (stress granule marker). Scale bar: 25 µm

## References

1. Philippe C, Burke S, Carrasco-Leon A, Martisova A, Casado P, Maniati E, et al. PERK orchestrates an endoplasmic reticulum stress alternative splicing program via CLK1/SRSF1. Nat Commun. 2026 Jun 19;17(1):7745. doi:10.1038/s41467-026-74397-y

2. Alberti S, Hyman AA. Biomolecular condensates at the nexus of cellular stress, protein aggregation disease and ageing. Nat Rev Mol Cell Biol. 2021 Mar;22(3):196–213. doi:10.1038/s41580-020-00326-6

3. Protter DSW, Parker R. Principles and Properties of Stress Granules. Trends in Cell Biology. 2016 Sep;26(9):668–79. doi:10.1016/j.tcb.2016.05.004

4. Nott TJ, Petsalaki E, Farber P, Jervis D, Fussner E, Plochowietz A, et al. Phase Transition of a Disordered Nuage Protein Generates Environmentally Responsive Membraneless Organelles. Molecular Cell. 2015 Mar;57(5):936–47. doi:10.1016/j.molcel.2015.01.013

5. Brangwynne CP, Eckmann CR, Courson DS, Rybarska A, Hoege C, Gharakhani J, et al. Germline P Granules Are Liquid Droplets That Localize by Controlled Dissolution/Condensation. Science. 2009 Jun 26;324(5935):1729–32. doi:10.1126/science.1172046

6. Namba H, Mir S, Groce EL, Alexander KA. Nuclear speckles: a fundamental layer of gene regulation. Trends in Cell Biology. 2026 Apr;S0962892426000590. doi:10.1016/j.tcb.2026.04.001

7. Lawrence IR, Sutton EC, Bhaskar S, Baserga SJ. Signaling to make human ribosomes: Connections between the cytoplasm and the nucleolus. Molecular Cell. 2026 Feb;86(3):469–90. doi:10.1016/j.molcel.2026.01.007

8. Holehouse AS, Alberti S. Molecular determinants of condensate composition. Mol Cell. 2025 Jan 16;85(2):290–308. doi:10.1016/j.molcel.2024.12.021 PubMed PMID: 39824169; PubMed Central PMCID: PMC11750178.

9. Greig JA, Nguyen TA, Lee M, Holehouse AS, Posey AE, Pappu RV, et al. Arginine-Enriched Mixed-Charge Domains Provide Cohesion for Nuclear Speckle Condensation. Molecular Cell. 2020 Mar;77(6):1237–1250.e4. doi:10.1016/j.molcel.2020.01.025

10. Molliex A, Temirov J, Lee J, Coughlin M, Kanagaraj AP, Kim HJ, et al. Phase separation by low complexity domains promotes stress granule assembly and drives pathological fibrillization. Cell. 2015 Sep 24;163(1):123–33. doi:10.1016/j.cell.2015.09.015 PubMed PMID: 26406374; PubMed Central PMCID: PMC5149108.

11. Ruff KM, King MR, Ying AW, Liu V, Pant A, Lieberman WE, et al. Molecular grammars of predicted intrinsically disordered regions that span the human proteome. Cell. 2026 Jan;189(1):323–342.e17. doi:10.1016/j.cell.2025.10.019

12. Lyons H, Veettil RT, Pradhan P, Fornero C, De La Cruz N, Ito K, et al. Functional partitioning of transcriptional regulators by patterned charge blocks. Cell. 2023 Jan 19;186(2):327–345.e28. doi:10.1016/j.cell.2022.12.013 PubMed PMID: 36603581; PubMed Central PMCID: PMC9910284.

13. Ngo JCK, Chakrabarti S, Ding JH, Velazquez-Dones A, Nolen B, Aubol BE, et al. Interplay between SRPK and Clk/Sty kinases in phosphorylation of the splicing factor ASF/SF2 is regulated by a docking motif in ASF/SF2. Mol Cell. 2005 Oct 7;20(1):77–89. doi:10.1016/j.molcel.2005.08.025 PubMed PMID: 16209947.

14. Masanta S, Wakimian K, Cieśla M. Phosphorylation-controlled nuclear speckle dynamics regulate splicing. Trends in Biochemical Sciences. 2026 Sep;51(9):896–909. doi:10.1016/j.tibs.2026.05.004

15. Guo YE, Manteiga JC, Henninger JE, Sabari BR, Dall’Agnese A, Hannett NM, et al. Pol II phosphorylation regulates a switch between transcriptional and splicing condensates. Nature. 2019 Aug;572(7770):543–8. doi:10.1038/s41586-019-1464-0 PubMed PMID: 31391587; PubMed Central PMCID: PMC6706314.

16. Zhang M, Gu Z, Sun Y, Dong Y, Chen J, Shu L, et al. Phosphorylation-dependent charge blocks regulate the relaxation of nuclear speckle networks. Mol Cell. 2025 May 1;85(9):1760–1774.e7. doi:10.1016/j.molcel.2025.03.016 PubMed PMID: 40233760.

17. McIntyre ABR, Tschan AB, Meyer K, Walser S, Rai AK, Fujita K, et al. Phosphorylation of a nuclear condensate regulates cohesion and mRNA retention. Nat Commun. 2025 Jan 4;16(1):390. doi:10.1038/s41467-024-55469-3

18. de Oliveira Freitas Machado C, Schafranek M, Brüggemann M, Hernández Cañás MC, Keller M, Di Liddo A, et al. Poison cassette exon splicing of SRSF6 regulates nuclear speckle dispersal and the response to hypoxia. Nucleic Acids Res. 2023 Jan ;51(2):870–90. doi:10.1093/nar/gkac1225 PubMed PMID: 36620874; PubMed Central PMCID: PMC9881134.

19. Jakubauskiene E, Vilys L, Makino Y, Poellinger L, Kanopka A. Increased Serine-Arginine (SR) Protein Phosphorylation Changes Pre-mRNA Splicing in Hypoxia. J Biol Chem. 2015 Jul 17;290(29):18079–89. doi:10.1074/jbc.M115.639690 PubMed PMID: 26023237; PubMed Central PMCID: PMC4505053.

20. Leva V, Giuliano S, Bardoni A, Camerini S, Crescenzi M, Lisa A, et al. Phosphorylation of SRSF1 is modulated by replicational stress. Nucleic Acids Res. 2012 Feb;40(3):1106–17. doi:10.1093/nar/gkr837 PubMed PMID: 21984412; PubMed Central PMCID: PMC3273819.

21. Becher I, Andrés-Pons A, Romanov N, Stein F, Schramm M, Baudin F, et al. Pervasive Protein Thermal Stability Variation during the Cell Cycle. Cell. 2018 May;173(6):1495–1507.e18. doi:10.1016/j.cell.2018.03.053

22. Sridharan S, Hernandez-Armendariz A, Kurzawa N, Potel CM, Memon D, Beltrao P, et al. Systematic discovery of biomolecular condensate-specific protein phosphorylation. Nat Chem Biol. 2022 Oct;18(10):1104–14. doi:10.1038/s41589-022-01062-y

23. Islam M, Rawnsley DR, Ma X, Navid W, Zhao C, Guan X, et al. Phosphorylation of CRYAB induces a condensatopathy to worsen post–myocardial infarction left ventricular remodeling. Journal of Clinical Investigation. 2025 Apr 1;135(7):e163730. doi:10.1172/JCI163730

24. Kuznetsova K, Scheremetjew M, Yin J, Moon H, Vargas DA, Hadarovich A, et al. CD-CODE 2.0: an enhanced condensate knowledgebase integrating pathobiology, condensate modulating drugs, and host–pathogen interactions. Nucleic Acids Research. 2026 Jan 654(D1):D375–82. doi:10.1093/nar/gkaf1104

25. Szklarczyk D, Kirsch R, Koutrouli M, Nastou K, Mehryary F, Hachilif R, et al. The STRING database in 2023: protein–protein association networks and functional enrichment analyses for any sequenced genome of interest. Nucleic Acids Research. 2023 Jan 6;51(D1):D638–46. doi:10.1093/nar/gkac1000

26. Paul S, Arias MA, Wen L, Liao SE, Zhang J, Wang X, et al. RNA molecules display distinctive organization at nuclear speckles. iScience. 2024 May;27(5):109603. doi:10.1016/j.isci.2024.109603

27. Decker CJ, Burke JM, Mulvaney PK, Parker R. RNA is required for the integrity of multiple nuclear and cytoplasmic membrane less RNP granules. EMBO J. 2022 May 2;41(9):EMBJ2021110137. doi:10.15252/embj.2021110137

28. Parker DM, Tauber D, Parker R. G3BP1 promotes intermolecular RNA-RNA interactions during RNA condensation. Molecular Cell. 2025 Feb;85(3):571–584.e7. doi:10.1016/j.molcel.2024.11.012

29. Hadarovich A, Singh HR, Ghosh S, Scheremetjew M, Rostam N, Hyman AA, et al. PICNIC accurately predicts condensate-forming proteins regardless of their structural disorder across organisms. Nat Commun. 2024 Dec 11;15(1):10668. doi:10.1038/s41467-024-55089-x PubMed PMID: 39663388; PubMed Central PMCID: PMC11634905.

30. Zhang M, Gu Z, Guo S, Sun Y, Ma S, Yang S, et al. SRRM2 phase separation drives assembly of nuclear speckle subcompartments. Cell Reports. 2024 Mar;43(3):113827. doi:10.1016/j.celrep.2024.113827

31. Okuda EK, Kessler LF, Arnold B, Riegger RJ, Hernández Cañás MC, Zebrowska E, et al. Rapid depletion and super-resolution microscopy reveal dual roles of SRSF5 in coordinating nuclear speckle–paraspeckle crosstalk during cellular stress. Nucleic Acids Research. 2025 Jul 19;53(14):gkaf713. doi:10.1093/nar/gkaf713

32. Tauber D, Tauber G, Khong A, Van Treeck B, Pelletier J, Parker R. Modulation of RNA Condensation by the DEAD-Box Protein eIF4A. Cell. 2020 Feb 6;180(3):411–426.e16. doi:10.1016/j.cell.2019.12.031 PubMed PMID: 31928844; PubMed Central PMCID: PMC7194247.

33. Maharana S, Wang J, Papadopoulos DK, Richter D, Pozniakovsky A, Poser I, et al. RNA buffers the phase separation behavior of prion-like RNA binding proteins. Science. 2018 May 25;360(6391):918–21. doi:10.1126/science.aar7366 PubMed PMID: 29650702; PubMed Central PMCID: PMC6091854.

34. Reis-Rodrigues P, Czerwieniec G, Peters TW, Evani US, Alavez S, Gaman EA, et al. Proteomic analysis of age-dependent changes in protein solubility identifies genes that modulate lifespan. Aging Cell. 2012 Feb;11(1):120–7. doi:10.1111/j.1474-9726.2011.00765.x PubMed PMID: 22103665; PubMed Central PMCID: PMC3437485.

35. Diner I, Nguyen T, Seyfried NT. Enrichment of Detergent-insoluble Protein Aggregates from Human Postmortem Brain. J Vis Exp. 2017 Oct 24;(128):55835. doi:10.3791/55835 PubMed PMID: 29155708; PubMed Central PMCID: PMC5755167.

36. Hadarovich A, Kuster D, Romero MLR, Toth-Petroczy A. On the Evolution of Biomolecular Condensates: From Prebiotic Origins to Subcellular Diversity. Annual Review of Cell and Developmental Biology. 2025 Oct 1;41(1):403–32. doi:10.1146/annurev-cellbio-101123-051723

37. Maharana S, Wang J, Papadopoulos DK, Richter D, Pozniakovsky A, Poser I, et al. RNA buffers the phase separation behavior of prion-like RNA binding proteins. Science. 2018 May 25;360(6391):918–21. doi:10.1126/science.aar7366 PubMed PMID: 29650702; PubMed Central PMCID: PMC6091854.

38. Parker DM, Tauber D, Parker R. G3BP1 promotes intermolecular RNA-RNA interactions during RNA condensation. Mol Cell. 2025 Feb 6;85(3):571–584.e7. doi:10.1016/j.molcel.2024.11.012 PubMed PMID: 39637853.

39. Markmiller S, Soltanieh S, Server KL, Mak R, Jin W, Fang MY, et al. Context-Dependent and Disease-Specific Diversity in Protein Interactions within Stress Granules. Cell. 2018 Jan 25;172(3):590–604.e13. doi:10.1016/j.cell.2017.12.032 PubMed PMID: 29373831; PubMed Central PMCID: PMC5969999.

40. Youn JY, Dunham WH, Hong SJ, Knight JDR, Bashkurov M, Chen GI, et al. High-Density Proximity Mapping Reveals the Subcellular Organization of mRNA-Associated Granules and Bodies. Molecular Cell. 2018 Feb;69(3):517–532.e11. doi:10.1016/j.molcel.2017.12.020

41. Reich S, Nguyen CDL, Has C, Steltgens S, Soni H, Coman C, et al. A multi-omics analysis reveals the unfolded protein response regulon and stress-induced resistance to folate-based antimetabolites. Nat Commun. 2020 Jun 10;11(1):2936. doi:10.1038/s41467-020-16747-y PubMed PMID: 32522993; PubMed Central PMCID: PMC7287054.

42. Ventura-Gomes A, Sousa-Luís R, Carmo-Fonseca M. Nuclear speckles: New insights into structure and function. Current Opinion in Structural Biology. 2026 Dec;101:103361. doi:10.1016/j.sbi.2026.103361

43. Wu J, Xiao Y, Liu Y, Wen L, Jin C, Liu S, et al. Dynamics of RNA localization to nuclear speckles are connected to splicing efficiency. Sci Adv. 2024 Oct 18;10(42):eadp7727. doi:10.1126/sciadv.adp7727

44. Williams TD, Michalak EM, Carey KT, Lam EYN, Anderson A, Griesbach E, et al. mRNA export factors store nascent transcripts within nuclear speckles as an adaptive response to transient global inhibition of transcription. Mol Cell. 2025 Jan 2;85(1):117–131.e7. doi:10.1016/j.molcel.2024.12.008 PubMed PMID: 39753105.

45. Dion W, Ballance H, Lee J, Pan Y, Irfan S, Edwards C, et al. Four-dimensional nuclear speckle phase separation dynamics regulate proteostasis. Sci Adv. 2022 Jan 7;8(1):eabl4150. doi:10.1126/sciadv.abl4150

46. Keshwani MM, Aubol BE, Fattet L, Ma CT, Qiu J, Jennings PA, et al. Conserved proline-directed phosphorylation regulates SR protein conformation and splicing function. Biochem J. 2015 Mar 1;466(2):311–22. doi:10.1042/BJ20141373 PubMed PMID: 25529026; PubMed Central PMCID: PMC5053020.

47. Ngo JCK, Chakrabarti S, Ding JH, Velazquez-Dones A, Nolen B, Aubol BE, et al. Interplay between SRPK and Clk/Sty Kinases in Phosphorylation of the Splicing Factor ASF/SF2 Is Regulated by a Docking Motif in ASF/SF2. Molecular Cell. 2005 Oct;20(1):77–89. doi:10.1016/j.molcel.2005.08.025

48. Ninomiya K, Adachi S, Natsume T, Iwakiri J, Terai G, Asai K, et al. LncRNA-dependent nuclear stress bodies promote intron retention through SR protein phosphorylation. EMBO J. 2020 Feb 3;39(3):e102729. doi:10.15252/embj.2019102729 PubMed PMID: 31782550; PubMed Central PMCID: PMC6996502.

49. Shinn MK, Tomares DT, Liu V, Pant A, Qiu Y, Vitalis A, et al. Nuclear speckle proteins form intrinsic and MALAT1-dependent microphases. Cell. 2026 Feb;189(3):832–852.e24. doi:10.1016/j.cell.2025.11.026

50. King MR, Ruff KM, Lin AZ, Pant A, Farag M, Lalmansingh JM, et al. Macromolecular condensation organizes nucleolar sub-phases to set up a pH gradient. Cell. 2024 Apr;187(8):1889–1906.e24. doi:10.1016/j.cell.2024.02.029

51. Jain S, Wheeler JR, Walters RW, Agrawal A, Barsic A, Parker R. ATPase-Modulated Stress Granules Contain a Diverse Proteome and Substructure. Cell. 2016 Jan 28;164(3):487–98. doi:10.1016/j.cell.2015.12.038 PubMed PMID: 26777405; PubMed Central PMCID: PMC4733397.

